# The *Braarudosphaera bigelowii* genome reveals cellular remodeling around a nitrogen-fixing organelle

**DOI:** 10.64898/2026.10.26.762046

**Authors:** Tyler H. Coale, Zoltán Füssy, Francisco M. Cornejo-Castillo, Wing Kwan Esther Mak, Kendra A. Turk-Kubo, Kyoko Hagino, Jonathan P. Zehr

## Abstract

*Braarudosphaera bigelowii* hosts the nitrogen-fixing cyanobacterial organelle UCYN-A, or the nitroplast, but the absence of a reference genome has limited investigation of the genomic features accompanying this association. Here, we present nuclear and organellar genome resources for the cultured *B. bigelowii* strain FR-21. The assembly comprises a 472.1-Mb nuclear genome together with representative nitroplast, chloroplast, and mitochondrial genomes of 1.49 Mb, 107 kb, and 44 kb, respectively. The four genomic compartments have strongly differentiated GC compositions, consistent with their distinct evolutionary origins.

The large nuclear genome does not appear to reflect a corresponding expansion of protein-coding capacity. Repeat annotation indicates that approximately 77% of the nuclear assembly is repetitive, leaving approximately 108 Mb of nonrepetitive sequence containing roughly 16,000 predicted protein-coding genes. Phylogenomic analysis places *B. bigelowii* within the expected diversity of haptophytes, indicating that its unusual diazotrophic lifestyle is not associated with an anomalous phylogenetic position.

Comparative analysis of nitrogen metabolism genes reveals extensive remodeling of host nitrogen acquisition and assimilation. *B. bigelowii* lacks major components required for nitrate and nitrite assimilation and has reduced capacity to use urea, cyanate, and several internally recycled nitrogen sources. In contrast, ammonium transport and the glutamine synthetase-glutamate synthase pathway are retained. This combination suggests reduced flexibility in nitrogen use and a host metabolism increasingly centered on ammonium supplied by the nitroplast.

*B. bigelowii* also encodes an expanded set of flavodoxin and cryptochrome genes. Eleven of 14 flavodoxin candidates and 21 of 39 cryptochrome candidates contained motifs linked to nitroplast targeting. Together with previous evidence that some of these proteins are abundantly imported into the nitroplast, these expansions point to host-encoded roles in nitroplast redox metabolism and diel regulation.

These genome resources establish a foundation for investigating nitroplast protein import, host-organelle metabolic integration, and the genomic consequences of acquiring an intracellular nitrogen-fixing compartment. They further show that the evolution of the *B. bigelowii*-UCYN-A relationship involved substantial remodeling of genome architecture and nitrogen metabolism in both partners.

## Introduction

Nitrogen availability constrains primary production across large regions of the ocean (Moore et al., 2001). Biological nitrogen fixation, the reduction of dinitrogen gas (N_2_) to ammonia, introduces new biologically available nitrogen into marine ecosystems and thereby influences ocean productivity and carbon cycling (Sohm et al., 2011; Zehr C Kudela, 2011). Although N_2_ fixation is catalyzed by prokaryote-derived nitrogenase, many environmentally important diazotrophs live in close association with photosynthetic eukaryotes (Foster C Zehr, 2019; Zehr C Capone, 2020). The repeated discovery of N_2_-fixing symbioses involving phylogenetically diverse marine phytoplankton suggests that these partnerships are not isolated anomalies, but a recurrent solution to nutrient limitation in the ocean. In such associations, photosynthetic hosts can provide fixed carbon and other resources that support the energetically demanding process of N_2_ fixation, while diazotrophs supply reduced nitrogen to hosts growing in nitrogen-limited waters. This reciprocal exchange links complementary metabolic capabilities and can enhance the growth and productivity of both partners(Foster et al., 2022). Such interactions are also evolutionarily important because prolonged metabolic dependence, cellular coordination, and genetic integration can move a partnership along a continuum from facultative association to obligate endosymbiosis and, ultimately, organellogenesis(Coale et al., 2024; Cornejo-Castillo et al., 2024).

As early molecular surveys began to reveal the diversity of N_2_-fixing microorganisms in the ocean, a distinct *nifH* lineage was repeatedly detected across geographically diverse marine environments and assigned the historical designation UCYN-A, for “unicellular cyanobacterial group A”(Church et al., 2005, 2008; Zehr et al., 1998). Subsequent surveys identified UCYN-A as one of the most widespread and abundant marine diazotrophic lineages, establishing its ecological importance before its cellular organization and association with eukaryotic hosts were understood(Church et al., 2008; Moisander et al., 2010). Subsequent genomic analyses revealed a highly reduced and metabolically unusual cyanobacterium. UCYN-A lacks oxygen-evolving photosystem II, the Calvin-Benson-Bassham cycle for carbon fixation, and the tricarboxylic acid cycle, while retaining photosystem I and the machinery required for N_2_ fixation (Tripp et al., 2010; Zehr et al., 2008). This combination distinguished UCYN-A from free-living cyanobacterial diazotrophs and suggested that it depended on an unidentified partner for fixed carbon and other essential metabolites.

Direct imaging and single-cell isotope analyses subsequently showed that UCYN-A associates with a unicellular haptophyte and transfers newly fixed nitrogen to its host in exchange for photosynthetically fixed carbon (Thompson et al., 2012). The larger coastal UCYN-A2 lineage was later identified within the haptophyte *Braarudosphaera bigelowii*, where the cyanobacterial partner corresponds to the intracellular structure historically described as the “spheroid body”(Hagino et al., 2013). *B. bigelowii* is a morphologically and genetically diverse haptophyte species complex that includes calcifying and non-calcifying life-cycle stages and has a fossil record extending into the Late Cretaceous (Hagino et al., 2013, 2026). Comparative analyses of UCYN-A1 and UCYN-A2 showed that the lineages share highly syntenic, reduced genomes and appear to have diverged after the major phase of genome reduction (Bombar et al., 2014). Environmental surveys further demonstrated strong fidelity between UCYN-A lineages and their respective haptophyte partners, consistent with a prolonged history of host-symbiont coevolution (Cornejo-Castillo et al., 2016).

The establishment of stable *B. bigelowii* cultures transformed this system from an environmentally observed association into an experimentally accessible model for N_2_-fixing endosymbiosis. Microscopy of cultured cells showed that UCYN-A2 is maintained intracellularly and divides in coordination with the host cell cycle (Suzuki et al., 2021). A host transcriptome also suggested that *B. bigelowii* expresses few of the pathways normally used by haptophytes to acquire alternative nitrogen sources. Transcripts encoding ammonium transport and assimilation were retained, whereas genes associated with nitrate, nitrite, urea, and cyanate utilization were not detected or were poorly represented (Suzuki et al., 2021). These observations raised the possibility that the host had become metabolically specialized around a dependable intracellular source of fixed nitrogen. However, because these conclusions were based primarily on transcriptomic data, it was not possible to determine whether undetected pathways were absent from the genome, transcriptionally silent under the conditions examined, expressed at levels below the detection threshold.

Additional evidence indicated that the relationship had progressed beyond a conventional obligate endosymbiosis. The relative sizes of UCYN-A and its haptophyte hosts are conserved across distinct symbiotic lineages and can be explained by metabolic trade-offs that coordinate growth and resource exchange between the partners(Cornejo-Castillo et al., 2024). Three-dimensional imaging of cultured *B. bigelowii* subsequently demonstrated that UCYN-A replication and division are integrated with the division of the host and its other organelles. Proteomic analysis also identified numerous proteins encoded by the algal nuclear genome but localized to UCYN-A, including proteins that supplement functions absent from the reduced cyanobacterial genome (Coale et al., 2024). Coordinated inheritance and the import of host-encoded proteins are defining characteristics of organellar integration. On this basis, UCYN-A was recognized as an early evolutionary-stage N_2_-fixing organelle, termed the nitroplast (Coale et al., 2024).

Recognition of the nitroplast changes the genomic unit that must be examined. The approximately 1.5-Mb UCYN-A genome represents only the genes retained within the organelle and cannot, by itself, describe the full functional repertoire of the nitroplast or the cellular changes supporting N_2_ fixation. Nitroplast biology also depends on genes encoded in the host nucleus, the targeting and import of their protein products, and the integration of N_2_ fixation with host carbon, energy, metal, and nitrogen metabolism. A complete host reference genome is therefore required to identify imported proteins systematically, distinguish gene loss from a lack of transcription, examine potential gene-family changes, and determine how the nuclear genome has been reorganized during the transition from symbiosis to organellogenesis.

Independent N_2_-fixing endosymbioses in other algae suggest that such transitions can produce extensive metabolic specialization in both partners. For example, *Epithemia* diatoms contain cyanobacterial endosymbionts that have lost broad metabolic autonomy and become specialized for intracellular N_2_ fixation (Moulin et al., 2024). Comparisons among these independently evolved systems offer an opportunity to identify common genomic consequences of acquiring an intracellular diazotroph. These consequences may include reductive evolution of the diazotroph, loss of redundant host nitrogen-acquisition pathways, altered protein-trafficking systems, and changes in host genome architecture.

Addressing these questions in *B. bigelowii*, however, has been limited by the absence of a nuclear reference genome. Previous work relied on environmental sequences, the reduced UCYN-A genome, or host transcriptomes, leaving nuclear genome size, repetitive content, protein-coding capacity, and comparative gene content unresolved.

Here, we present a nuclear and organellar genome resource for the cultured *B. bigelowii* strain FR-21, which contains a UCYN-A2 nitroplast. We characterize the nuclear, nitroplast, chloroplast, and mitochondrial genomic compartments and compare their sizes and nucleotide compositions. We place *B. bigelowii* within a phylogenomic and comparative genomic framework of available haptophyte genomes, evaluate the relationship between nuclear genome size and protein-coding capacity, and examine the contribution of repetitive sequence to its expanded nuclear genome. We additionally screened the complete nuclear protein-model set for nitroplast-associated and conventional N-terminal targeting signals and tested whether flavodoxin and cryptochrome/photolyase families had expanded in *B. bigelowii* relative to other haptophytes. Finally, we use the genome to test previous predictions concerning host nitrogen metabolism and to identify genomic losses and retained pathways associated with ammonium acquisition and assimilation. These resources establish a genomic foundation for studying nitroplast protein import, host-organelle metabolic integration, and the broader evolutionary consequences of acquiring a N_2_-fixing organelle.

## Results

### Assembly of the B. bigelowii nuclear and organellar genomes

Long-read sequencing of *B. bigelowii* FR-21 produced assemblies representing the host nuclear genome and its three genetically distinct organelles: the nitroplast, chloroplast, and mitochondrion. The curated nuclear assembly comprised 472,100,991 bp distributed across 587 sequences, with an overall GC content of 60.24% (Supplementary Dataset S1). The representative nitroplast, chloroplast, and mitochondrial genomes were 1,493,781 bp, 107,247 bp, and 44,277 bp, respectively (Fig. 1A). The nuclear assembly had an N50 of 1,475,824 bp, reached with 97 contigs (Supplementary Fig. S1). Combined PacBio HiFi and Oxford Nanopore mean sequencing depth averaged 58.3x across nuclear contigs, with each contig weighted equally (Supplementary Fig. S2).

**Figure 1.**
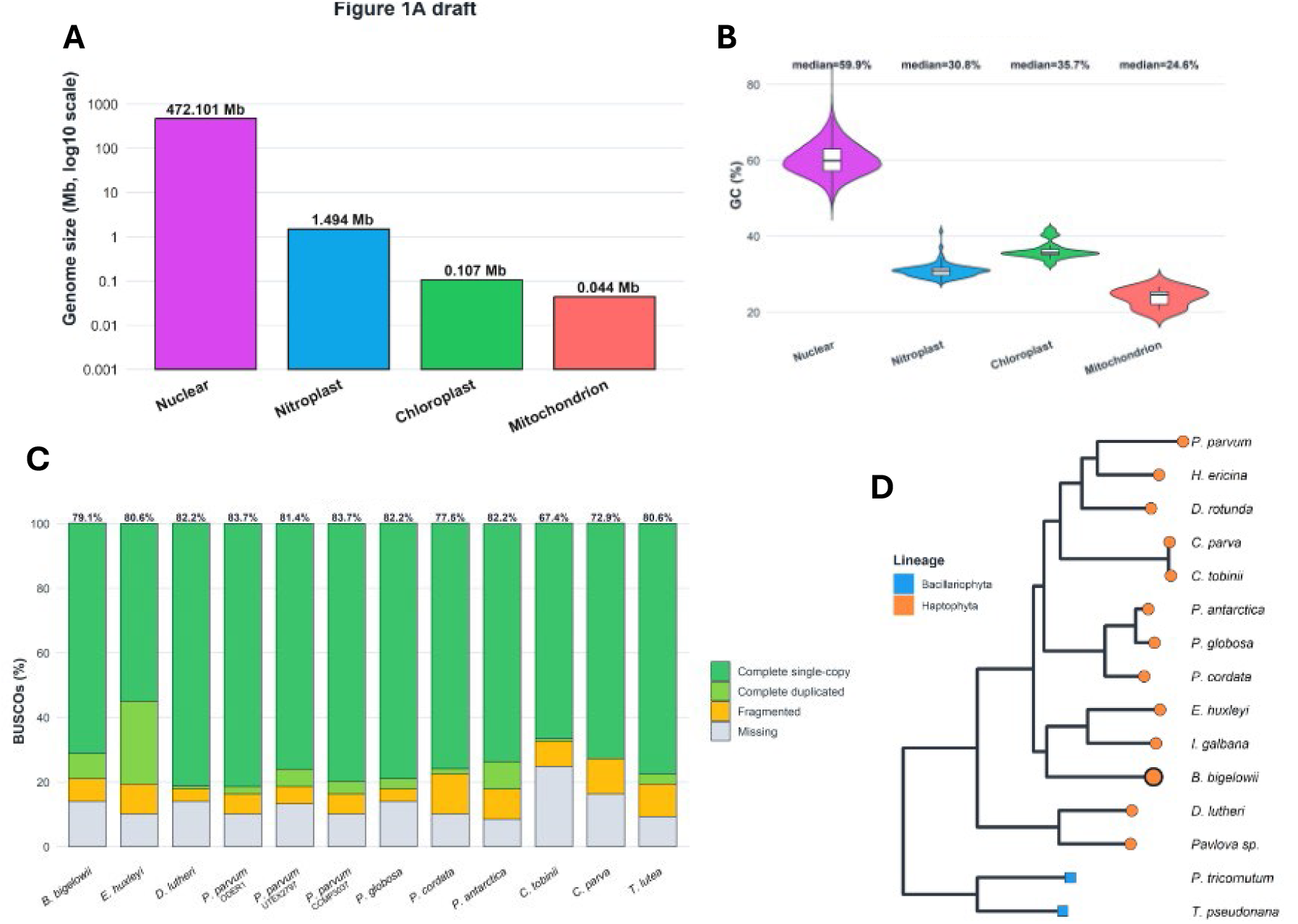
Genome architecture, assembly completeness, and phylogenomic placement of *B. bigelowii* FR-21. (A) Sizes of the nuclear, nitroplast, chloroplast, and mitochondrial genomes. Genome sizes are plotted on a log10 scale and shown above each bar. (B) GC-content distributions calculated in 10-kb windows with a 5-kb step across the four genomic compartments. Violin plots show the distribution of windowed GC content, with internal boxplots indicating the median and interquartile range; median values are reported above each distribution. Only canonical A, C, G, and T bases were included when calculating GC percentage. (C) BUSCO assessment of *B. bigelowii* and 11 comparison haptophyte genome assemblies using the eukaryota_odb12 dataset of 129 conserved orthologs. Bars show the percentages of complete single-copy, complete duplicated, fragmented, and missing BUSCOs. Values above the bars indicate the total percentage of complete BUSCOs. The same BUSCO workflow and lineage dataset were applied to all assemblies. (D) Rooted species tree inferred from the predicted proteomes of *B. bigelowii*, 12 comparison haptophytes, and the diatoms *Phaeodactylum tricornutum* and *Thalassiosira pseudonana* using OrthoFinder. Orange circles denote haptophytes and blue squares denote the diatom outgroups; *B. bigelowii* is highlighted with a black outline. Nuclear assembly contiguity and combined long-read coverage are shown in Supplementary Figs. S1 and S2, respectively. Contig-classification evidence and comparative genome resources are provided in Supplementary Datasets S1 and S2.

The four genomic compartments differed markedly in nucleotide composition. GC content calculated in 10-kb sliding windows had median values of 59.9% for the nuclear genome, 30.8% for the nitroplast, 35.7% for the chloroplast, and 24.6% for the mitochondrial genome (Fig. 1B). The nuclear genome therefore had substantially higher GC content than all three organellar genomes, whereas the mitochondrial genome had the lowest GC content. The distinct distributions, rather than only differences in genome-wide means, provided clear compositional separation among the four compartments.

Assembly completeness was evaluated using BUSCO with the eukaryota_odb12 dataset (Manni et al., 2021). The *B. bigelowii* assembly contained complete matches to 79.1% of the 129 markers. This value was within the range observed for the comparison haptophyte assemblies, which recovered between 67.4% and 83.7% complete BUSCOs when analyzed using the same lineage dataset and gene-prediction workflow (Fig. 1C). These results support the use of the FR-21 assembly for downstream comparative analyses of conserved gene content and genome evolution.

### The nuclear genome is phylogenetically consistent with established haptophyte relationships

To place the new genome in a comparative framework, a species tree was inferred from curated proteomes representing 13 haptophytes, with the diatoms *Phaeodactylum tricornutum and Thalassiosira pseudonana* included as outgroups (Supplementary Dataset S2). Orthogroups and the rooted species tree were inferred using OrthoFinder (Emms C Kelly, 2019).

The resulting phylogeny recovered the expected separation of haptophytes from the diatom outgroups and grouped closely related haptophyte taxa together (Fig. 1D). The two *Chrysochromulina* species formed a clade, the three *Phaeocystis* species grouped together, and *Diacronema lutheri* grouped with the other pavlovalean representative. These relationships were broadly concordant with published haptophyte genome studies (Füssy et al., 2025; Hulatt et al., 2021; Read et al., 2013; Wisecaver et al., 2023).

*B. bigelowii* was placed within the principal haptophyte radiation, adjacent to the lineage containing *Emiliania huxleyi* and *Isochrysis galbana* (Fig. 1D). Its placement did not indicate an unusually long or isolated lineage relative to other sampled haptophytes. The distinctive genomic features of FR-21 therefore occur within an otherwise recognizable haptophyte phylogenomic context rather than reflecting an anomalous taxonomic placement.

### Repeat expansion accounts for most of the large nuclear genome

The 472.1-Mb *B. bigelowii* nuclear assembly was substantially larger than most of the available haptophyte genomes included in the comparison (Fig. 2A; Supplementary Dataset S2). To determine whether this difference reflected increased protein-coding capacity or expansion of repetitive sequence, repeats were identified de novo and used to mask the nuclear assembly(Flynn et al., 2020).

**Figure 2.**
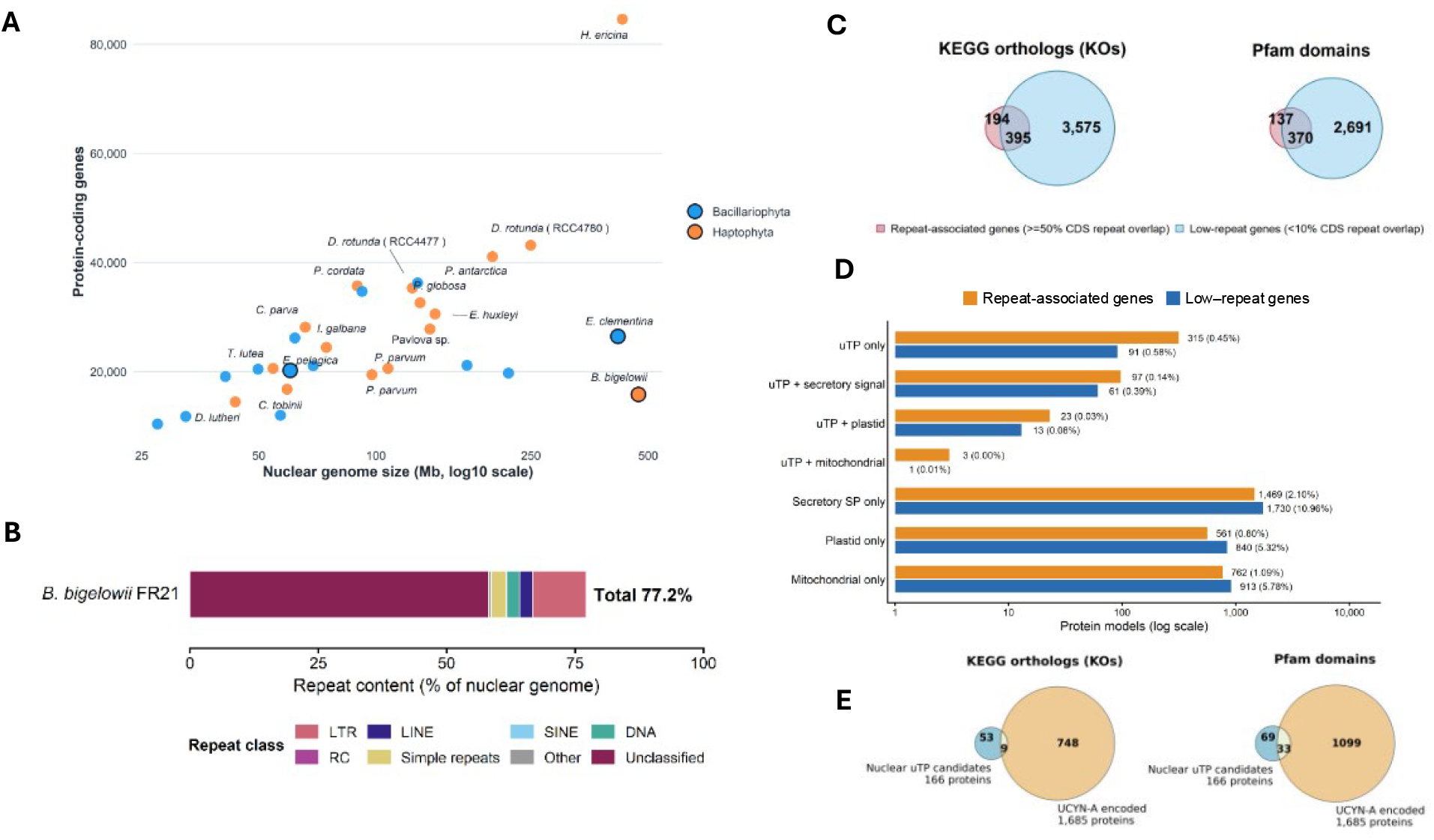
Repeat expansion underlies the large *B. bigelowii* nuclear genome. (A) Relationship between nuclear genome size and predicted protein-coding gene number in *B. bigelowii* and comparison algal genomes. Genome size is shown on a log10 scale. Orange and blue points denote haptophytes and diatoms, respectively. For *B. bigelowii*, the protein-coding gene count represents the conservative set of 15,788 nuclear models with less than 10% CDS overlap with annotated repeats and no detected transposable-element domain. (B) Repeat composition of the *B. bigelowii* FR-21 nuclear genome. Repeats were identified using a species-specific RepeatModeler library and classified with RepeatMasker. The stacked bar shows the percentage of the nuclear assembly assigned to long terminal repeat retrotransposons (LTRs), long and short interspersed nuclear elements (LINEs and SINEs), DNA transposons, rolling-circle elements (RC), simple repeats, other repeat classes, and unclassified repeats. (C) Overlap of functional annotations identified among repeat-associated genes, defined as having at least 50% CDS overlap with annotated repeats, and low-repeat genes, defined as having less than 10% CDS repeat overlap. Venn diagrams show unique and shared KEGG Orthology identifiers and Pfam domains. Circle areas are proportional to the total number of distinct annotations in each gene set; numbers indicate annotations unique to either set or shared between them. (D) Predicted targeting-signals among all 85,753 nuclear protein models. Rows are mutually exclusive combinations of uTP motif evidence, a TargetP 2.0 non-plastid secretory-signal prediction, an ASAFind 2.0 high- or low-confidence plastid prediction, and a TargetP 2.0 mitochondrial-transit-peptide prediction. Bars show protein-model counts on a logarithmic scale; labels report counts and percentages within each gene-model class (<10% CDS-repeat overlap and no detected TE-domain evidence versus all remaining gene models). (E) Venn diagrams compare the distinct KEGG Orthology identifiers and Pfam domains assigned to the 166 nuclear proteins in the low-repeat set that met the uTP criterion and the 1,685 proteins encoded by the UCYN-A2 genome. Values are numbers of distinct annotation terms rather than proteins. Repeat and gene-filtering classifications are provided in Supplementary Dataset S3, and targeting predictions and UCYN-A annotation-overlap counts in Supplementary Dataset S4. The corresponding overlap comparisons for all nuclear uTP candidates and the low-repeat subset are shown in Supplementary Fig. S3.

A total of 364,402,329 bp, corresponding to 77.19% of the nuclear assembly, was classified as repetitive. The remaining unmasked portion comprised 107,698,662 bp, or 22.81% of the assembly (Fig. 2B; Supplementary Dataset S3). Unclassified repeats were the largest category, accounting for 58.15% of the nuclear assembly. Long terminal repeat retrotransposons represented a further 10.43%, while DNA transposons and LINE elements accounted for 2.53% and 2.54%, respectively. SINE elements were rare, representing approximately 0.004% of the assembly. Thus, both classified transposable elements and a large pool of lineage-specific or otherwise unclassified repetitive sequences contributed to nuclear genome expansion.

We predicted 15,788 nuclear protein-coding genes in non-repetitive regions of the *B. bigelowii* genome. This gene set was defined by low coding-sequence overlap with annotated repeats (<10% of CDS length) and no detectable transposable element domains, and we use it as a conservative estimate of the conventional host gene repertoire (Supplementary Dataset S3 and S11). Despite the large size of the nuclear assembly, this estimate places *B. bigelowii* within the range of protein-coding gene counts observed for other haptophyte genomes (Fig. 2A).

We also recovered a much larger set of predicted coding sequences within repeat-rich regions of the genome. These repeat-associated predictions accounted for most of the raw gene-model total, but they were functionally annotated much less often than genes in non-repetitive regions. Approximately 65.0% of low-repeat models received an eggNOG annotation, compared with 29.6% of repeat-rich models. Nevertheless, comparisons of annotation terms showed that the repeat-associated set was not devoid of recognizable functions and included many annotations not found in the low-repeat gene set (Fig. 2C).

These unique annotations likely reflect, at least in part, the biology of transposable elements, repeat-associated coding sequences, and other gene predictions embedded in repetitive regions, rather than a proportional expansion of the conventional host gene repertoire.

To assess the broader contribution of host nuclear genes to organelle biology, we screened all 85,753 nuclear protein models for candidate UCYN-A (uTP), secretory, plastid, and mitochondrial targeting signals and summarized their co-occurrence (Fig. 2D;

Supplementary Dataset S4). The low-repeat set contained 3,649 of 15,788 models with at least one form of targeting evidence, compared with 3,230 of the 69,965 models excluded by the repeat/TE filter. In total, 604 proteins met the uTP criterion. Of these, 406 lacked a detected N-terminal targeting signal, 158 had a non-plastid secretory prediction, 36 had a plastid-targeting prediction, and four had a mitochondrial-targeting prediction. Among proteins without uTP evidence, 3,199 were classified as non-plastid secretory candidates, 1,401 as plastid-targeting candidates, and 1,675 as mitochondrial-targeting candidates; 78,874 lacked all four signals. Each targeting category represented a higher proportion of the low-repeat set than of the excluded models. To determine whether candidate host imports primarily duplicate functions already encoded by UCYN-A or expand its functional repertoire, we compared annotations assigned to the 166 uTP-positive proteins in the low-repeat set with those assigned to the 1,685 UCYN-A2-encoded proteins. Only 9 of the 62 distinct KEGG Orthology identifiers detected among the nuclear candidates were also detected among UCYN-A proteins, leaving 53 (85.5%) unique to the import candidate set. Similarly, only 33 of 102 Pfam domains were shared, whereas 69 (67.6%) occurred only among the import candidates. Thus, most functional annotations associated with the candidate imported proteins were not represented in the UCYN-A-encoded proteome (Fig. 2E; Supplementary Dataset S4). The expanded comparison includes all 604 nuclear uTP candidates and the 166 candidates retained in the low-repeat gene set, each compared with the UCYN-A-encoded proteome (Supplementary Fig. S3).

### Chloroplast architecture is more conserved than mitochondrial architecture among haptophytes

Whole-genome alignments revealed contrasting patterns of structural conservation in the *B. bigelowii* chloroplast and mitochondrial genomes relative to other haptophytes (Fig. 3; Supplementary Dataset S5). The chloroplast genomes retained broad collinearity across the six representative haptophytes shown in the main-text comparison (Fig. 3A).

**Figure 3.**
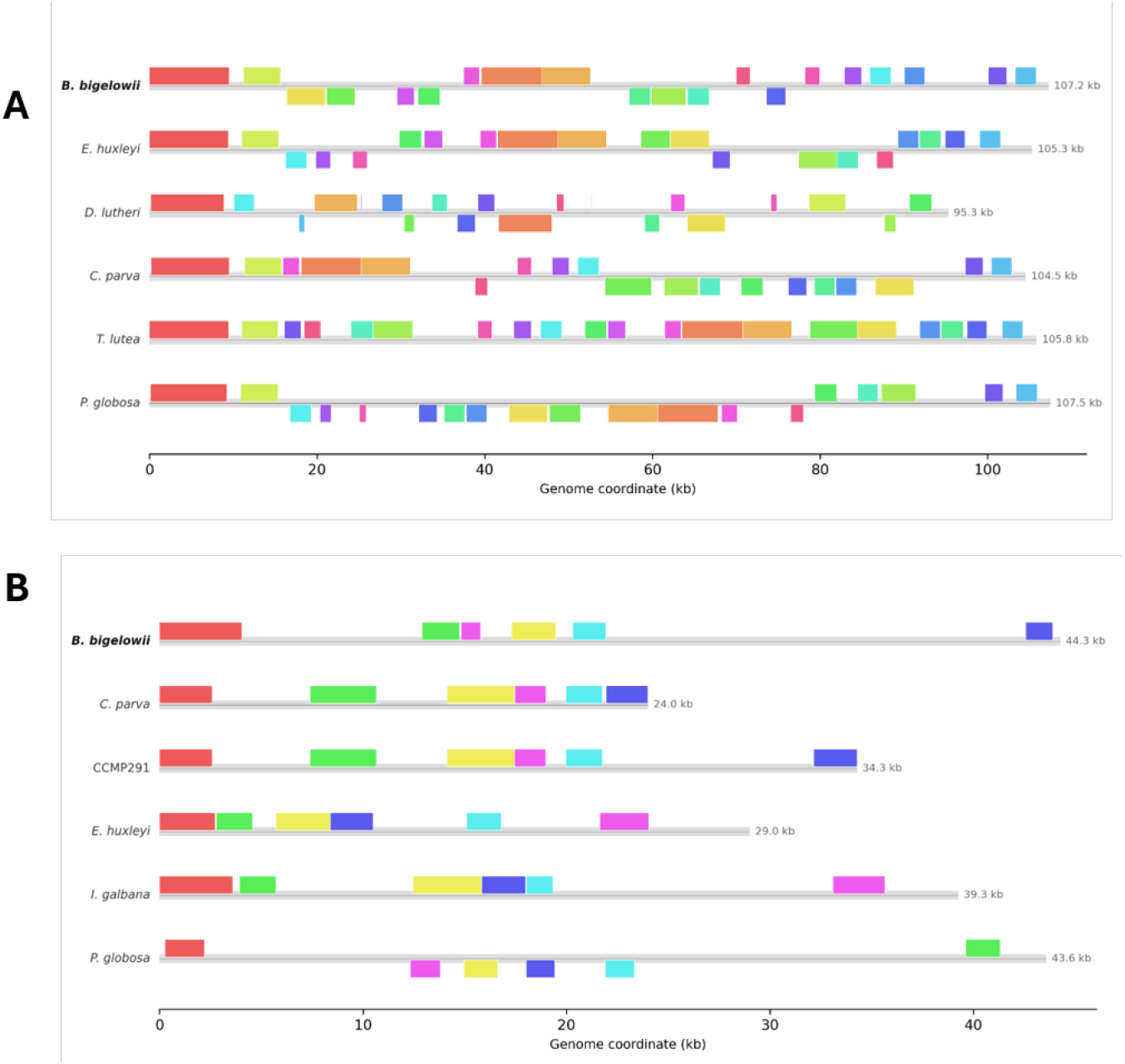
Chloroplast genome architecture is more conserved than mitochondrial genome architecture among haptophytes. (A) Whole-genome alignment of the *B. bigelowii* FR-21 chloroplast genome with chloroplast genomes from *Emiliania huxleyi*, *Diacronema lutheri*, *Chrysochromulina parva*, *Tisochrysis lutea*, and *Phaeocystis globosa*. (B) Whole-genome alignment of the *B. bigelowii* mitochondrial genome with mitochondrial genomes from *C. parva*, *Chrysochromulina* sp. CCMP291, *E. huxleyi*, *Isochrysis galbana*, and *P. globosa*. Circular genomes were initially aligned with progressiveMauve, rotated to a common locally collinear block (LCB), reverse-complemented when required to place the anchor in the same orientation, and then realigned. Colored rectangles represent homologous LCBs present in all six genomes with a median span of at least 1 kb; matching colors identify corresponding blocks among genomes. Blocks plotted below the genome centerline are in reverse orientation relative to the rotated reference. Gray bars represent complete genome sequences, with genome lengths shown at right. Uncolored regions were not retained under the shared-block filtering criteria. Expanded chloroplast and mitochondrial comparisons are shown in Supplementary Fig. S4A and S4B, respectively; genome accessions and comparison metadata are provided in Supplementary Dataset S5.

After filtering to locally collinear blocks (LCBs) present in all six taxa and with a median length of at least 1 kb, 20 shared blocks were retained. Together, these blocks spanned 63.6% of the 107,247-bp *B. bigelowii* chloroplast genome. Most occurred in similar order and orientation among taxa, although several inversions and local rearrangements were evident. The expanded nine-taxon comparison showed the same overall pattern while recovering additional smaller and less broadly shared regions (Supplementary Fig. S4A).

The mitochondrial alignment was substantially more fragmented (Fig. 3B). Only six LCBs were present in all six representative taxa and had a median length of at least 1 kb, together spanning 27.1% of the 44,277-bp *B. bigelowii* mitochondrial genome. Shared mitochondrial regions varied more extensively in order and orientation and were separated by larger lineage-specific or unaligned intervals. This pattern was also evident in the expanded seven-genome comparison (Supplementary Fig. S4B). These alignments indicate greater conservation of chloroplast genome architecture than mitochondrial genome architecture among the haptophytes examined. *B. bigelowii* followed this general pattern, with broadly conserved chloroplast organization and more variable mitochondrial organization, and showed no conspicuous departure from the structural variation observed among the comparison taxa.

### The B. bigelowii genome has lost major nitrogen assimilation pathways

Nitrogen-metabolism proteins were identified and manually curated across *B. bigelowii* and 12 comparison haptophytes (Fig. 4; Supplementary Dataset S6). The analysis included proteins involved in nitrate and nitrite assimilation, ammonium transport and assimilation, urea and cyanate utilization, and arginine metabolism.

**Figure 4.**
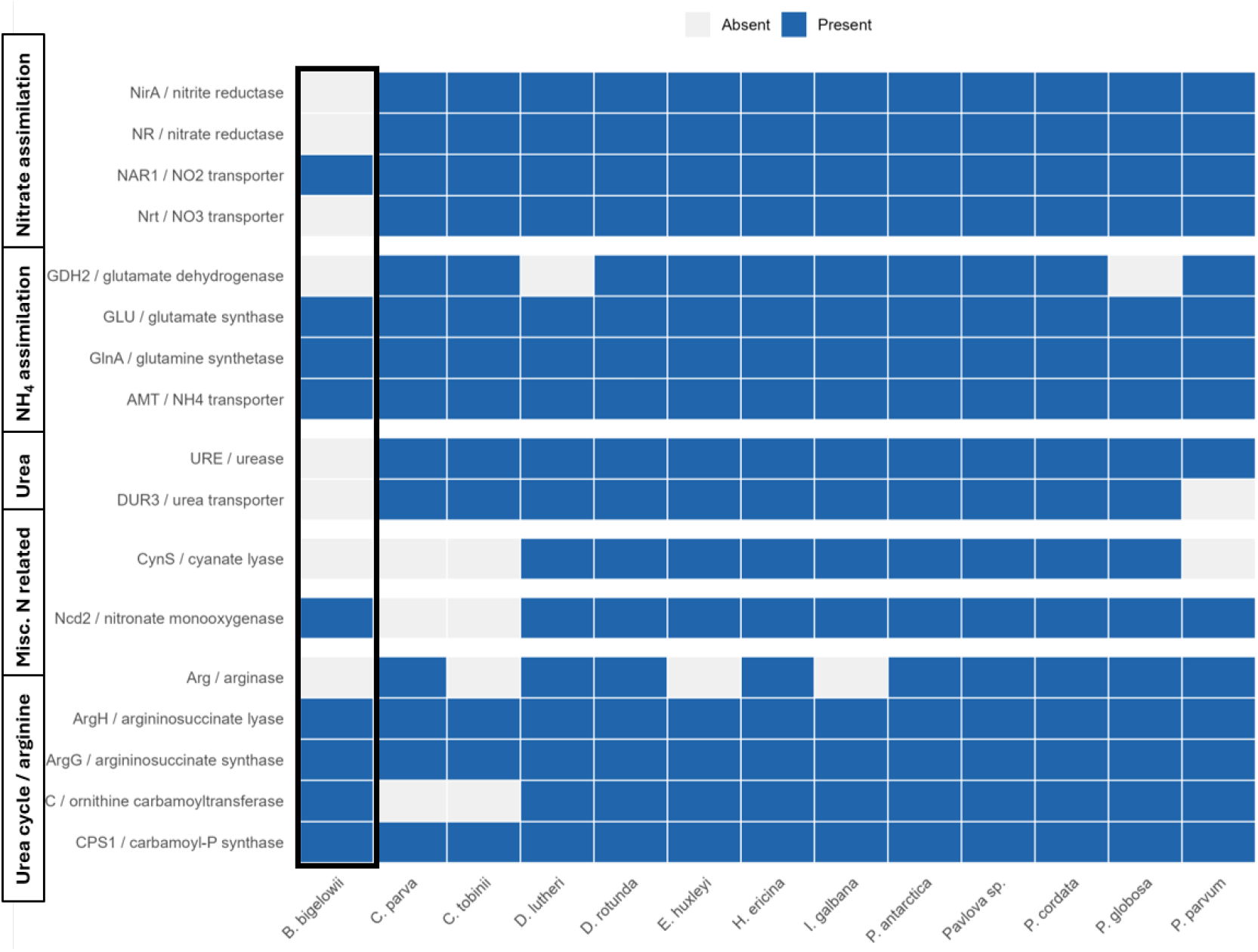
Remodeling of host nitrogen acquisition and assimilation pathways in *B. bigelowii*. Presence-absence matrix of nitrogen-metabolism protein families in *B. bigelowii* and 12 comparison haptophytes. Candidate homologs were identified from standardized eggNOG-mapper annotations and manually reviewed; a protein family was scored as present when at least one supported homolog was retained. Blue and gray cells indicate presence and absence, respectively, and the black outline highlights *B. bigelowii*. For *B. bigelowii*, only host nuclear-encoded proteins were considered; nitroplast, chloroplast, and mitochondrion proteins were excluded. Curated marker-family counts and supporting annotation evidence are provided in Supplementary Dataset S6.

*B. bigelowii* lacked the principal components of the canonical assimilatory nitrate pathway. No homologs of the major nitrate transporter, nitrate reductase, or assimilatory nitrite reductase were recovered. Each of these components was detected in all other haptophytes included in the comparison (Fig. 4). Homologs of the nitrate assimilation-related protein / formate–nitrite transporter family were present in *B. bigelowii*. The retention of these genes is difficult to interpret in isolation because this transporter family includes proteins with different substrate specificities, including nitrite, bicarbonate, formate, and related anions. These homologs may therefore transport substrates other than nitrite, represent vestiges of a previously complete nitrate-assimilation pathway, or have roles in nitrite handling that are not yet understood. However, in the absence of nitrate reductase and assimilatory nitrite reductase, these proteins do not constitute a complete pathway for reducing environmental nitrate or nitrite to ammonium. The genomic results therefore support the conclusion that FR-21 lacks conventional nitrate assimilation rather than merely failing to express the pathway under previously examined culture conditions.

Capacity for urea utilization was also reduced. No urease homolog was detected in the *B. bigelowii* gene set, and no clearly assigned degradation of urea (DUR)-family urea transporter was recovered. Urease was present in every other haptophyte examined (Fig. 4), indicating that loss of the enzyme required to break down urea is a distinctive feature of *B. bigelowii* within this comparison. Urea-transport annotations were more variable across taxa: for example, *P. parvum* lacked a DUR-family homolog but contained proteins annotated with urea channel activity. The absence of urease in *B. bigelowii*, together with the lack of a clearly identified canonical urea transporter, indicates that urea is unlikely to serve as a direct nitrogen source through the conventional urease pathway.

Cyanate lyase was also absent from *B. bigelowii*. Because this enzyme was missing from several other haptophytes as well, its absence was not unique to FR-21, but it reduces the set of nitrogen substrates for which the host genome encodes a recognizable utilization pathway. In contrast, two nitronate monooxygenase-like proteins were retained (Fig. 4; Supplementary Dataset S6). These proteins may participate in nitronate or nitroalkane metabolism and could release nitrite from organic nitronate compounds. However, because no assimilatory nitrite reductase was recovered, this pathway does not provide a clear route for incorporating nitronate-derived nitrite into ammonium.

*B. bigelowii* retained genes for ammonium transport and assimilation, including the core machinery needed to incorporate reduced nitrogen. The nuclear genome encodes 19 AMT-family ammonium-transporter homologs, three glutamine synthetase proteins, and three glutamate synthase proteins. These genes retain the GS–GOGAT pathway for incorporating ammonium into glutamine and glutamate.

No glutamate dehydrogenase (GDH) homolog was recovered from the *B. bigelowii* genome (Fig. 4). Although GDH was also absent from several comparison haptophytes and is therefore not uniquely missing from the nitroplast-bearing lineage, its absence leaves GS-GOGAT as the only recognized route for net ammonium assimilation into glutamate.

This pattern is consistent with a reduced repertoire of enzymes linking ammonium and glutamate metabolism relative to haptophytes that encode both GS-GOGAT and GDH.

Most of the canonical arginine-biosynthesis pathway was retained. The *B. bigelowii* genome encodes five carbamoyl-phosphate synthase homologs and single homologs of ornithine carbamoyltransferase, argininosuccinate synthase, and argininosuccinate lyase. No arginase homolog was recovered (Fig. 4; Supplementary Dataset S6). Thus, the genome retains the downstream enzymatic steps for arginine synthesis but lacks the conventional arginine-catabolic reaction that converts arginine to ornithine and urea. Arginase was also absent from several comparison haptophytes and therefore does not represent a lineage-specific loss in *B. bigelowii*.

### B. bigelowii encodes an expanded set of ffavodoxins and cryptochrome/photolyases with uTP-associated motifs

Broad screening of eggNOG annotations recovered 14 nuclear-encoded flavodoxin candidates from *B. bigelowii* and 36 candidates across the 12 comparison haptophyte proteomes (Fig. 5A; Supplementary Dataset S7). *B. bigelowii* contained more candidates than any comparison genome, which encoded between zero and eight. Ten *B. bigelowii* models belonged to the low-repeat set, whereas four had ≥10% overlap with repeat regions and were retained as additional candidates.

**Figure 5.**
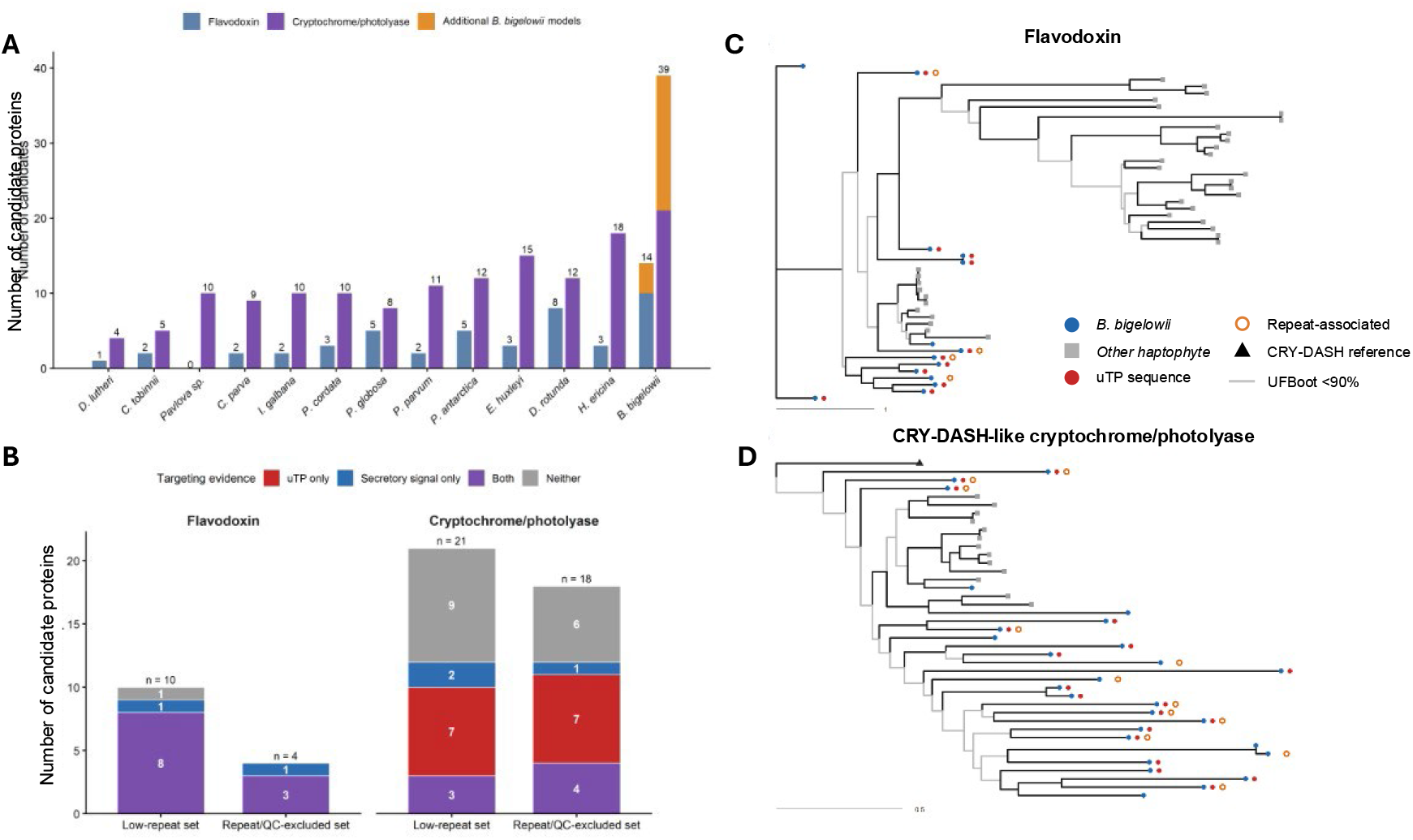
Expansion and putative nitroplast targeting of flavodoxin and cryptochrome/photolyase proteins in *B. bigelowii*. (A) Paired bars show the numbers of flavodoxin (blue) and cryptochrome/photolyase-family (purple) candidate protein models recovered from the *B. bigelowii* nuclear genome and 12 comparison haptophyte proteomes; values above the bars give the candidate count for each family. For *B. bigelowii*, orange portions show additional models excluded by the repeat/ǪC filter: four of the 14 flavodoxins and 18 of the 39 cryptochrome/photolyase proteins. The remaining portions of the *B. bigelowii* bars represent the 10 flavodoxin and 21 cryptochrome/photolyase models retained in the low-repeat nuclear gene set. (B) Targeting-signal evidence among *B. bigelowii* flavodoxin and cryptochrome/photolyase candidates, partitioned by protein family and CDS overlap with annotated repeats. Colors indicate proteins with significant terminal uTP-associated motif evidence only, a TargetP 2.0 secretory-signal prediction only, both signals, or neither signal. Numbers within segments give candidate counts, and labels above bars give total sample sizes. (C) Maximum-likelihood tree of 14 *B. bigelowii* and 36 comparison-haptophyte flavodoxin candidates, inferred under Ǫ.PFAM+F+G4 from the aligned N-terminal 210 amino acids of each protein, using the complete sequence when shorter. (D) CRY-DASH-associated subtree of the maximum-likelihood cryptochrome/photolyase phylogeny, containing the *Synechocystis* CRY-DASH reference P77967 and all 40 proteins in its haptophyte sister lineage. The phylogeny was inferred under LG+F+R4 from 489 positions of the conserved photolyase-homology region defined by P77967, excluding variable terminal extensions. In both (C) and (D) panels, all branches are expanded and tip labels are omitted for clarity. Blue circles identify *B. bigelowii* proteins, gray squares identify proteins from other haptophytes, and the black triangle in (D) identifies P77967. Red circles denote terminal uTP motif evidence, and open orange circles mark *B. bigelowii* models with ≥10% CDS overlap with annotated repeats. Branches leading to internal nodes with <90% ultrafast-bootstrap support are light gray; other branches are black. Support was assessed with 1,000 ultrafast-bootstrap replicates, numerical support values are provided in Supplementary Figs. S6 and S8, and branch lengths represent substitutions per site. Full-length motif maps are shown in Supplementary Figs. S5 and S7. Candidate annotations are provided in Supplementary Datasets S7 and S8, protein sequences in Supplementary Dataset S9, and CDS sequences in Supplementary Dataset S10.

Eleven of the 14 *B. bigelowii* flavodoxin candidates contained significant C-terminal uTP-associated motif matches, whereas none of the 36 flavodoxins identified in the 12 comparison haptophytes met the same criterion (Fig. 5B; Supplementary Fig. S5). The three *B. bigelowii* exceptions were g4905.t1, g75613.t1 and g89161.t1. Thus, uTP-associated sequence features were strongly enriched among the expanded *B. bigelowii* flavodoxin set.

The maximum-likelihood tree placed the *B. bigelowii* candidates in several distinct parts of the haptophyte flavodoxin tree rather than in a single species-specific cluster (Fig. 5C; Supplementary Fig. S6). Based on their eggNOG orthogroup assignments, five *B. bigelowii* candidates were classified as bacterial and nine as eukaryotic. Three of the five bacterial-classified *B. bigelowii* candidates also contained uTP-associated motifs. Five of the 14 *B. bigelowii* candidates (35.7%) were classified as bacterial, compared with four of 36 candidates (11.1%) across the comparison haptophytes; *B. bigelowii* therefore contained the largest absolute number of bacterial-classified flavodoxins and a 3.2-fold higher proportion than the pooled comparison set. Only one bacterial-classified flavodoxin (g39669.t1) received a cyanobacterial assignment (Supplementary Fig. S6; Supplementary Dataset S7). However, blastp searches against the NCBI ClusteredNR database recovered haptophyte proteins as the closest matches, followed by numerous cyanobacterial and other bacterial homologs, indicating that this protein is broadly conserved among haptophytes and may derive from a cyanobacterial ancestor rather than representing a recent endosymbiotic gene transfer from UCYN-A.

Of the three flavodoxin candidates lacking sufficient uTP evidence, one (g4905.t1) was repeat-associated, with 50.9% of its CDS overlapping annotated repeats, while another (g75613.t1) contained two uTP-associated motifs but did not meet the three-motif threshold (Supplementary Fig. S5; Supplementary Dataset S7). Thus, only one flavodoxin gene model (g89161.t1) unequivocally lacked evidence of nitroplast targeting.

The same search identified 39 cryptochrome/photolyase candidates in *B. bigelowii* and 124 across the 12 comparison haptophytes, with four to 18 candidates in each comparison proteome (Fig. 5A; Supplementary Dataset S8). 21 *B. bigelowii* candidates belonged to the low-repeat set and 18 had ≥10% CDS-repeat overlap. A separate 21 of the 39 *B. bigelowii* candidates met the uTP detection criterion, whereas none of the comparison-haptophyte candidates did (Fig. 5B; Supplementary Fig. S7). uTP-positive proteins occurred in both gene sets: 10 of 21 low-repeat and 11 of 18 repeat-excluded candidates.

Phylogenetic analysis placed 40 haptophyte proteins in a CRY-DASH-like lineage sister to the curated *Synechocystis* sp. PCC 6803 CRY-DASH protein P77967 (97% ultrafast-bootstrap support for the sister relationship; Fig. 5D; Supplementary Fig. S8). This lineage contained 27 *B. bigelowii* proteins and 13 representatives collectively spanning all 12 comparison haptophytes. 19 of the 27 *B. bigelowii* proteins, but none of the 13 comparison proteins, contained uTP sequences (Fig. 5D; Supplementary Fig. S7). 7 *B. bigelowii* members of the lineage received cyanobacterial eggNOG assignments (Supplementary Fig. S8; Supplementary Dataset S8). All 40 lineage members were closer to the CRY-DASH reference than to the seven other curated cryptochrome and photolyase references included in the analysis. These results support a *B. bigelowii*-specific expansion within an ancestrally haptophyte CRY-DASH-like family, but do not resolve whether its members function primarily as photolyases, photoreceptors, or other flavoproteins.

## Discussion

### Host-genome remodeling around nitroplast-derived nitrogen

The *B. bigelowii* genome provides a genomic basis for an unusual feature of the UCYN-A symbiosis that has emerged from both culture and field studies: the host appears to rely heavily on nitrogen supplied by its N_2_-fixing partner even when alternative dissolved nitrogen sources are available. NanoSIMS measurements of natural UCYN-A symbioses showed little or no host assimilation of nitrate and relatively little contribution of external ammonium to host nitrogen demand, with N_2_ fixation by UCYN-A instead supplying a substantial fraction of host nitrogen requirements even under dissolved-inorganic-nitrogen-replete conditions (Mills et al., 2020). Transcriptomic analysis of cultured *B. bigelowii* similarly failed to detect expression of many genes required for the uptake and assimilation of nitrogen sources other than ammonium, although the absence of transcripts could not distinguish gene loss from condition-dependent expression (Suzuki et al., 2021). The genome resolves much of this uncertainty. *B. bigelowii* lacks nitrate reductase and ferredoxin-dependent nitrite reductase, as well as conventional nitrate transporters, and therefore does not encode a complete pathway for reducing environmental nitrate or nitrite to ammonium. It also lacks urease and DUR3-family urea transporters, while the absence of cyanate lyase, glutamate dehydrogenase, and arginase further reduces alternative routes for nitrogen utilization and intracellular recycling (Fig. 4).

In contrast, the machinery required to assimilate reduced nitrogen is retained. The genome encodes AMT-family ammonium transporter homologs, glutamine synthetases, and glutamate synthases, preserving the GS-GOGAT pathway that incorporates ammonium into glutamine and glutamate (Fig. 4; Supplementary Dataset S6). The pattern of nitrogen-related gene loss reflects a narrowing of the routes available for nitrogen acquisition and recycling, while retaining the machinery required to assimilate reduced nitrogen. The large AMT repertoire is particularly intriguing in this context. Earlier transcriptomic work proposed ammonium transport as a potential component of nitrogen exchange between UCYN-A and its host, but the cellular localization and substrate specificity of the individual *B. bigelowii* AMTs remain unknown. Their expansion therefore should not be taken as evidence for high environmental ammonium uptake; instead, some family members may participate in intracellular nitrogen trafficking associated with the nitroplast. Determining the localization and transport properties of these proteins will be important for resolving how fixed nitrogen moves from the nitroplast into host metabolism.

Loss of alternative nitrogen-utilization pathways may also help explain the ecological behavior of *Braarudosphaera*-UCYN-A associations. UCYN-A is not restricted to oligotrophic waters depleted in nitrogen and can be an important contributor to N_2_ fixation in coastal and otherwise nutrient-rich environments (Cabello et al., 2020; Fletcher-Hoppe et al., 2023; Muñoz-Marín et al., 2019). *Braarudosphaera*-UCYN-A associations, for example, can account for much of the measured N_2_ fixation in portions of the Southern California Current System(Turk-Kubo et al., 2021). If the host cannot directly use environmental nitrate, nitrate availability would not reduce its need for internally supplied fixed nitrogen and would therefore not be expected to suppress N_2_ fixation, consistent with observations from cultured *B. bigelowii* (Coale et al., 2024). The genome thus provides a metabolic basis for the occurrence of *Braarudosphaera*-UCYN-A associations in environments where external nitrogen sources such as nitrate are available. *B. bigelowii*, however, is not completely dependent on the nitroplast for all nitrogen acquisition.

Phagotrophy has been suggested as an additional potential route for acquisition of organic nutrients, and the relative importance of this pathway in natural populations remains unresolved (Mak et al., 2024).

Whether loss of nitrogen-metabolism pathways was itself selectively advantageous is less clear. Once nitroplast-derived reduced nitrogen became a reliable resource, pathways for nitrate, urea, and other nitrogen sources may simply have experienced relaxed selection. Their subsequent loss could nevertheless reinforce host dependence on the nitroplast by reducing the metabolic alternatives available if nitrogen fixation is lost.

Maintenance of redundant nitrogen-acquisition pathways also carries energetic and regulatory costs, raising the possibility that pathway loss became advantageous after the symbiosis was established. These alternatives cannot presently be distinguished, but both imply progressive remodeling of host metabolism following sustained access to fixed nitrogen. Such losses may therefore represent part of the transition from a metabolically complementary symbiosis toward a more obligate cellular integration.

The expanded *B. bigelowii* nuclear genome, at 472 Mb, is substantially larger than many sequenced haptophyte genomes and primarily reflects accumulation of repetitive DNA rather than a proportional expansion of the conventional protein-coding gene repertoire. Approximately 77% of the assembly is repetitive, including a large unclassified fraction and substantial LTR-retrotransposon content, whereas the conservative low-repeat gene set contains approximately 16,000 protein-coding genes (Fig. 2A,B; Supplementary Dataset S3). A similar decoupling of genome size from protein-coding capacity has recently been reported for the endosymbiont-bearing diatom *Epithemia clementina*, in which genome expansion is also dominated by repetitive sequence (Frail et al., 2025). However, this pattern is not universal among algae with N_2_-fixing endosymbionts: the related marine diatom *E. pelagica* lacks comparable repeat expansion despite harboring a diazotrophic endosymbiont(Frail et al., 2025; Schvarcz et al., 2022). This variation suggests that repeat expansion is not an inevitable consequence of endosymbiosis or organelle acquisition. The photosynthetic chromatophore of *Paulinella* species provides the only other known recent example of primary endosymbiosis and shows that organelle integration, gene transfer, protein import, and host-genome remodeling can occur without a single predictable pattern of nuclear genome expansion (Lhee et al., 2021; Nowack et al., 2008; Singer et al., 2017; Stephens et al., 2021).

Nevertheless, repeat proliferation could provide a permissive genomic environment for the evolutionary remodeling that accompanies endosymbiotic integration. An instructive precedent comes from the independently evolved chromatophore of *P. micropora*, where retrotransposition generated additional copies of endosymbiont-derived nuclear genes and facilitated their regulatory rewiring during establishment of the photosynthetic organelle (Calatrava et al., 2022). Transposable elements and other repetitive sequences can promote gene duplication, rearrangement, and regulatory innovation, potentially increasing the genomic substrate on which selection can act as a host accommodates a newly integrated compartment (Chuong et al., 2017; Feschotte, 2008). The present data do not establish that repeat expansion in *B. bigelowii* was caused by, or directly facilitated, nitroplast evolution. However, the combination of extensive repeat accumulation, pronounced remodeling of nitrogen metabolism, and the emergence of a host-controlled N_2_-fixing organelle raises the possibility that host-genome restructuring and metabolic specialization proceeded together during the evolution of the *B. bigelowii*-UCYN-A association.

### Toward nitrogen-fixing eukaryotes

Engineering N_2_ fixation into a photosynthetic eukaryote has long been a goal of plant synthetic biology, but the problem extends well beyond expression of the nitrogenase structural proteins. Nitrogenase requires a suite of accessory proteins for metallocluster assembly and maturation, substantial inputs of ATP and reducing power, and protection from oxygen (Burén et al., 2020; Dixon C Kahn, 2004). Considerable progress has been made toward reconstructing these requirements experimentally. For example, refactoring of the *Cyanothece* N_2_-fixation cluster in the oxygenic phototroph *Synechocystis* demonstrated that a defined set of nif and accessory genes can support nitrogenase activity, while also showing that the relative expression of these components strongly affects activity (Liu et al., 2025). In parallel, approaches for reconstructing nitrogenase in eukaryotic organelles have demonstrated stable, stoichiometric expression and processing of multiple Nif proteins targeted to yeast mitochondria, providing a potential strategy for coordinating the many components required for nitrogenase assembly(Yang et al., 2023). Nevertheless, the complete cellular machinery required for N_2_ fixation in an aerobic photosynthetic cell remains unresolved.

The nitroplast provides a complementary view of this engineering problem: it represents a naturally evolved solution in which N_2_ fixation has already been integrated into an oxygenic photosynthetic eukaryote. In *B. bigelowii*, oxygen evolution is confined to the chloroplast, whereas N_2_ fixation occurs in a separate intracellular compartment that is metabolically and structurally integrated with the host. The nitroplast imports host-encoded proteins and coordinates its division with the host cell, demonstrating that stable N_2_ fixation in a eukaryote involves not only nitrogenase but also extensive exchange of metabolites, proteins, and regulatory information across the host-organelle interface (Coale et al., 2024).

A synthetic nitrogen-fixing compartment would require more than nitrogenase activity alone; it would also need efficient metabolic exchange with the host. Fixed nitrogen must be exported from the diazotrophic compartment and incorporated into host metabolism, while the host must provide substrates sufficient to meet the substantial carbon, energy, and reductant demands of nitrogen fixation. Carbon import and fixed-nitrogen export have similarly been identified as key unresolved interfaces in evaluating N_2_-fixing endosymbionts as potential engineering platforms (Elhai, 2026).

*E. clementina* provides a useful comparison because genomic and proteomic analyses suggest that its strategy may rely on host-derived carbon to support ATP and reductant generation while also consuming oxygen, potentially creating conditions favorable for nitrogenase activity during host photosynthesis (Moulin et al., 2024). Similar questions remain unresolved for the nitroplast. The identity of the carbon substrates supplied by *B. bigelowii*, the sources of reductant and ATP supporting nitrogen fixation, and the mechanisms that protect nitrogenase from oxygen during daytime N_2_ fixation are not yet known.

Flavodoxin provides one candidate mechanism for supplying reductant to the nitroplast. Reduced UCYN-A genomes lack flavodoxin, indicating that this electron-transfer protein was lost during genome reduction, although they retain several ferredoxins that have been proposed to supply electrons to nitrogenase (Bombar et al., 2014; Tripp et al., 2010). In the related free-living diazotroph *Crocosphaera watsonii*, however, flavodoxin accumulates together with nitrogenase during the dark period even under iron-replete conditions, consistent with preferential use of flavodoxin rather than ferredoxin for electron transfer to nitrogenase (Saito et al., 2011). Previous proteomic analysis of the nitroplast identified a host-encoded flavodoxin containing a uTP among the most abundant *B. bigelowii* proteins detected in UCYN-A during the day, raising the possibility that the host has replaced a redox function lost from the UCYN-A genome (Coale et al., 2024). The expanded flavodoxin repertoire identified here strengthens this hypothesis: 11 of the 14 nuclear *B. bigelowii* flavodoxin candidates contain uTP-associated motifs (Fig. 5A,B; Supplementary Fig. S5). These proteins could potentially couple reducing equivalents generated by host metabolism to redox reactions within the nitroplast, including nitrogenase reduction. Whether flavodoxin is reduced before import, receives electrons after entering the nitroplast, or participates in another redox-transfer pathway is unknown, and these paralogs may not be functionally equivalent. The close metabolic and physical association of the nitroplast with the chloroplast and mitochondrion further raises the possibility that host-encoded flavodoxins participate in redox coupling among these compartments, although there is presently no direct evidence for such a mechanism.

The loss of flavodoxin from the nitroplast genome also has implications for iron metabolism, as it is one of the typical mechanisms for iron conservation in marine phytoplankton (La Roche et al., 1996). The nitroplast has lost access to exogenous iron sources and lost iron uptake transporters (Cheung et al., 2026; Tripp et al., 2010), and relies on iron acquired by the haptophyte uptake machinery including phytotransferrin and ISIP3 (Behnke C LaRoche, 2020; Lampe et al., 2024). The impact of a new, iron-rich organelle on the whole-cell iron budget, as well as the details of intracellular iron trafficking remain unknown.

The cryptochrome/photolyase family shows an even larger expansion. *B. bigelowii* encodes 39 candidates, compared with 4 to 18 in each comparison haptophyte, and 21 of the 39 candidates had uTP motifs (Fig. 5A,B; Supplementary Fig. S7). Most of the expansion falls within a CRY-DASH-like lineage represented in all 12 comparison haptophytes, consistent with expansion of an existing haptophyte family rather than acquisition of an entirely novel protein class (Fig. 5D; Supplementary Fig. S8). This lineage was sister to the manually reviewed UniProtKB/Swiss-Prot CRY-DASH protein P77967 from *Synechocystis* sp. PCC 6803, and 19 of its 27 *B. bigelowii* members were nitroplast-targeted (Fig. 5D; Supplementary Figs. S7 and S8). CRY-DASH proteins retain the photolyase-homology region and have been associated with single-stranded nucleic-acid repair and light-responsive regulatory functions (Kleine et al., 2003; Rredhi et al., 2021; Selby C Sancar, 2006). The expansion of putatively nitroplast-targeted cryptochrome/photolyases therefore suggests a role in light-dependent regulation of nitroplast activity. Cryptochrome-like proteins occur in cyanobacteria, where they are evolutionarily and functionally distinct from DNA-repair photolyases (Hitomi et al., 2000), and nuclear-encoded cryptochromes can also be targeted to chloroplasts, where they regulate photosynthetic and other plastidial processes (Rredhi et al., 2021, 2024). The *B. bigelowii* expansion may therefore represent a host-encoded regulatory system that couples nitroplast physiology to light availability and the diel cycle. Direct localization, spatial proteomics, diel expression measurements, and functional characterization will be needed to distinguish these possibilities.

The limited functional overlap between the conservative uTP candidate set and the UCYN-A proteome suggests that host protein import expands the functional repertoire of the nitroplast rather than simply duplicating functions that remain encoded by UCYN-A (Fig. 2E; Supplementary Fig. S3; Supplementary Dataset S4). Candidate imports may therefore compensate for functions lost during UCYN-A genome reduction while also introducing host-derived regulatory, repair, transport, and metabolic capabilities not encoded by the present UCYN-A genome. Because this comparison is based on predicted uTP motifs and broad functional annotations, it does not demonstrate nitroplast localization or biochemical function for individual proteins. Nevertheless, the predominance of import candidate-specific annotations suggests that host protein import has expanded the functional repertoire of the nitroplast in parallel with reduction of the UCYN-A genome.

The genome-wide targeting screen suggests that host contributions to nitroplast biology may extend beyond the protein families examined here. Most of the 604 uTP-containing proteins lacked a conventional N-terminal targeting prediction, consistent with a targeting signal that is distinct from standard secretory, plastid, or mitochondrial presequences. Nevertheless, 198 uTP-positive proteins also carried an N-terminal prediction, most commonly a secretory signal (Fig. 2D; Supplementary Dataset S4). This overlap could reflect genuine dual or multi-step targeting, but could also arise from ambiguous sequence features, prediction errors, or non-functional remnants of ancestral targeting sequences. The present analysis cannot distinguish among these possibilities.

The high degree of integration in *B. bigelowii* may make the UCYN-A nitroplast difficult to transfer directly into a new host. UCYN-A has lost extensive biosynthetic and metabolic capacity, and some of these losses are complemented by host-encoded proteins imported into the nitroplast, while others likely require metabolite or nutrient exchange with the host. A new host would therefore need to reproduce these dependencies or replace the missing functions through other means. More broadly, the portability of all known nitroplasts and diazotrophic symbionts remains uncertain, and each is likely to depend on additional proteins, metabolic exchanges, and regulatory mechanisms that have not yet been identified. Elhai (2026) similarly argued that the extensive host dependence of UCYN-A makes it a less tractable starting point for transplantation than more metabolically autonomous N_2_-fixing endosymbionts. At the same time, this high degree of integration may be especially informative for engineering because it provides an opportunity to identify host-derived functions that have evolved to couple eukaryotic and diazotrophic metabolism. Such functions could reveal mechanisms that increase N_2_ fixation efficiency or rates, improve exchange between the N_2_-fixing compartment and its host, or otherwise enhance the performance of an engineered system.

Together, these emerging diazotrophic systems provide natural examples for identifying which features are necessary for establishing N_2_ fixation in a eukaryotic cell and which arise through deeper integration over evolutionary time. Comparative analysis of these systems may distinguish a minimal set of functions required to establish a stable N_2_-fixing compartment from additional host-derived proteins and regulatory mechanisms that improve its performance. The *B. bigelowii* genome, combined with nitroplast proteomics, now makes it possible to identify these host contributions directly and to test whether selected eukaryotic proteins could enhance N_2_ fixation when introduced into a cyanobacterial chassis. In this way, the nitroplast offers not a ready-made solution for transplantation, but an evolved blueprint for discovering new components and design principles for engineering N_2_-fixing eukaryotes.

## Methods

### B. bigelowii culture, DNA sequencing, and genome assembly

*B. bigelowii* strain FR-21, which contains the UCYN-A2 nitroplast (Hagino et al., 2026), was cultured in modified f/2 seawater media with no added nitrate, as described in Coale et al. (2024). Cultures were maintained at 20 °C under a 12 h light:12 h dark cycle at an irradiance of 40 µmol photons m^-2^ s^-1^. Cells were harvested during exponential growth for genomic DNA extraction.

High-molecular-weight (HMW) genomic DNA was extracted at the University of California, Davis from approximately 20,000,000 cells collected as cell pellets and flash frozen in liquid nitrogen. DNA extraction was performed using a freeze–thaw, SDS– proteinase K, and S-CTAB protocol. The resulting DNA had a 260/280 ratio of 1.83, a 260/230 ratio of 2.21, a concentration of 79 ng/µL, and a total yield of approximately 7 µg (88 µL total volume).

HMW DNA was used for PacBio HiFi library preparation and sequencing on one SMRT cell at UC Davis. PacBio HiFi reads were assembled using hifiasm v0.25.0 with default parameters, including a 51-bp k-mer, three rounds of haplotype-aware error correction, four rounds of assembly-graph cleaning, and default haplotig purging. Read-coverage thresholds used during assembly and haplotig purging were inferred automatically by hifiasm. This generated primary and alternate pseudohaplotype assemblies.

In parallel, Oxford Nanopore Technologies (ONT) libraries were prepared from the same HMW DNA. Additional long-read sequencing was performed at the UC Davis DNA Technologies Core using an ONT PromethION platform. Libraries were prepared using the Ligation Sequencing Kit V14 and sequenced on an R10.4.1 PromethION flow cell using standard sequencing without adaptive sampling. Sequence data were basecalled using the super-accuracy model in the ONT basecalling software implemented through MinKNOW. The run generated 8.5 Gb of sequence data with a read N50 of 14.5 kb. ONT reads were used as an independent long-read dataset to support assembly validation and polishing.

Assembly contigs were manually classified as nuclear, bacterial, nitroplast, chloroplast, or mitochondrial using multiple independent lines of evidence (Supplementary Dataset S1). Taxonomic assignments were initially assessed using Kraken2 v2.1.3 with a confidence threshold of 0.1 (--confidence 0.1) and otherwise default parameters, and sequencing reads were mapped to the assembly to evaluate contig coverage and coverage consistency. Predicted proteins were searched against the reviewed UniProtKB/Swiss-Prot database (release 2025_04) using DIAMOND BLASTP v2.2.5 in sensitive mode with the BLOSUM62 scoring matrix and otherwise default parameters.

The highest-scoring Swiss-Prot match for each protein was retained and assigned to its corresponding taxonomic domain (Eukaryota, Bacteria, Archaea, or viruses), and the distribution of these assignments was summarized for each contig. Classification was further informed by contig GC content and the presence of compartment-specific marker proteins identified from protein-sequence similarity searches. Chloroplast contigs were identified by photosynthesis-associated marker proteins, including Rbc and Psb proteins, mitochondrial contigs by Cob and Cox proteins, and nitroplast contigs by NifH protein similarity to published UCYN-A2 NifH. Final compartment assignments were made by integrating taxonomic, read-coverage, GC-content, and marker-protein evidence (Supplementary Dataset S1).

Contigs classified as nuclear in the curated contig-classification table were retained to define the final nuclear assembly. This yielded 587 nuclear contigs totaling 472,100,991 bp. The complete evidence underlying the individual contig assignments is provided in Supplementary Dataset S1.

Representative nitroplast, chloroplast, and mitochondrial sequences were manually curated separately from the nuclear genome. The final organelle representatives corresponded to contig_13101 (nitroplast), contig_12873 (chloroplast), and ptg001320l (mitochondrion). These curated organelle sequences replaced earlier fragmented bins in all downstream analyses.

### Assembly quality assessment

Assembly continuity statistics, including Nx values, were calculated directly from contig lengths in the final curated nuclear genome assembly. Contigs were ranked from longest to shortest, and Nx was defined as the contig length at which the cumulative assembly length reached x% of the total assembly length (Supplementary Fig. S1). Long-read support for the final assembly was evaluated independently using the PacBio HiFi and ONT datasets. Reads were aligned to the curated nuclear genome using minimap2 v2.30. PacBio HiFi reads were aligned using the HiFi preset (-ax map-hifi), corresponding to the minimap2 high-accuracy long-read settings (-k19 -w19 -U50,500 -g10k -A1 -B4 -O6,26 - E2,1 -s200). ONT reads were aligned using the ONT long-read preset (-ax map-ont). All other minimap2 parameters were left at their defaults.

Alignments were converted to BAM format, coordinate-sorted, and indexed using SAMtools v1.22 with default parameters. Per-contig sequencing support was summarized using samtools coverage with default parameters. Coverage was defined as the percentage of reference positions with at least one aligned base, and mean sequencing depth was calculated for each contig. Under the SAMtools v1.22 defaults, no minimum mapping- or base-quality threshold was imposed, and unmapped, secondary, ǪC-failed, and duplicate-marked alignments were excluded from coverage calculations. For Supplementary Fig. S2, the PacBio HiFi and ONT mean depths were summed for each contig, and the reported mean was calculated across contigs with equal weight per contig.

Genome completeness was evaluated using BUSCO v6.0.0 with the eukaryota_odb12 lineage dataset in genome mode and Miniprot v0.18 as the gene predictor, with otherwise default parameters. The eukaryota_odb12 dataset contained 129 conserved single-copy orthologs. The same BUSCO workflow and lineage dataset were applied to *B. bigelowii* and all comparison haptophyte genome assemblies to permit direct comparison of assembly completeness (Fig. 1C).

GC-content distributions were calculated directly from the curated nuclear and organelle genome sequences. GC content was measured in 10-kb windows with a 5-kb step. Only canonical A, C, G, and T bases were included in the denominator when calculating GC percentage; ambiguous bases were excluded (Fig. 1B).

### Nuclear and organelle gene prediction

Nuclear protein-coding genes were predicted using AUGUSTUS v3.5.0 with a *B. bigelowii*-specific training model and otherwise default prediction parameters. The species-specific model was trained using the assembled *B. bigelowii* genome together with transcript evidence from an rnaSPAdes assembly of previously generated FR-21 RNA-seq data (Suzuki et al., 2021) after removal of low-coverage and non-eukaryotic transcripts.

Coding sequences corresponding to the final curated nuclear assembly were reconstructed from the AUGUSTUS annotations after restricting predictions to the 587 contigs classified as nuclear. This yielded 85,753 nuclear AUGUSTUS gene models before repeat filtering.

Organelle gene annotation was treated separately from nuclear gene prediction. The initial AUGUSTUS annotation did not recover genes from the prokaryotic-like nitroplast genome; in particular, the principal nitroplast contig (contig_13101) contained no AUGUSTUS gene models despite strong sequence evidence identifying it as UCYN-A. Organelle genomes were subsequently manually curated and circularized, and curated chloroplast and mitochondrial protein-coding annotations were generated separately from the nuclear AUGUSTUS annotation.

For the final nitroplast gene set, protein-coding sequences were predicted directly from the curated contig_13101 sequence using Prodigal v2.6.3 with genetic code 11 (Bacterial, Archaeal and Plant Plastid Code) and otherwise default parameters. This yielded 1,685 nitroplast CDS predictions. All predicted CDS had lengths divisible by three and lacked internal stop codons under translation table 11. The final genome annotation therefore combined AUGUSTUS predictions for the nuclear genome, curated chloroplast and mitochondrial annotations, and Prodigal predictions for the nitroplast.

### Repeat identification and repeat gene filtering

Repetitive sequences in the nuclear assembly were identified using a de novo repeat-discovery and masking workflow. A species-specific repeat library was generated using RepeatModeler v2.0.7 with RMBlast v2.14.1 as the search engine (Flynn et al., 2020). A RepeatModeler database was first generated from the curated nuclear genome using BuildDatabase, followed by repeat-family discovery using 12 threads and otherwise default parameters. The optional LTR structural-discovery module (-LTRStruct) was not enabled.

The resulting classified repeat-family consensus library was used for genome-wide repeat annotation with RepeatMasker v4.2.3 using RMBlast and the parameters -pa 12 -engine rmblast -xsmall -gff -lib, where -lib specified the *B. bigelowii*-specific RepeatModeler library. RepeatMasker therefore produced both repeat annotations and a soft-masked nuclear genome. Repeat annotations were summarized by major repeat class, including long terminal repeat (LTR) retrotransposons, LINEs, SINEs, DNA transposons, and unclassified repeats (Fig. 2B; Supplementary Dataset S3). For the stacked repeat-composition plot, each repeat-covered nuclear base was assigned to the overlapping RepeatMasker hit with the highest alignment score, yielding mutually exclusive class totals; percentages were calculated relative to the 472,100,991-bp nuclear assembly.

Because the initial AUGUSTUS annotation contained a large number of predicted genes within repeat-rich regions, repeat overlap was quantified for each nuclear gene model. Repeat intervals were parsed from the RepeatMasker .out file, overlapping repeat intervals were merged to avoid double-counting, and the number of CDS bases overlapping annotated repeats was calculated for each AUGUSTUS model. CDS repeat-overlap fraction was calculated as the number of CDS bases overlapping one or more RepeatMasker annotations divided by the total CDS length of the gene model. Models with less than 10% CDS overlap with annotated repeats were classified as low-repeat gene models.

Predicted proteins were also screened using InterProScan v5.76-107.0 with default parameters to identify protein-domain evidence characteristic of transposable elements. Repeat-overlap and protein-domain evidence were integrated into a repeat/TE quality-control classification. The most conservative nuclear gene set consisted of models assigned to the class low_repeat_no_TE_domain_hit, defined by less than 10% CDS overlap with annotated repeats and no detected InterProScan transposable-element domain evidence. This procedure retained 15,788 nuclear AUGUSTUS CDS and was used as the conservative estimate of the conventional protein-coding gene repertoire. The complete set of 85,753 AUGUSTUS predictions was retained separately to avoid discarding potentially functional genes solely because of their genomic context (Supplementary Dataset S3).

### Genome-wide prediction of protein-targeting signals

All 85,753 predicted *B. bigelowii* nuclear proteins were screened for uTP-associated motifs and conventional secretory, plastid, and mitochondrial targeting signals. uTP-associated motifs were detected with MAST from MEME Suite v5.5.9 (Bailey et al., 2009, 2015), using the motif model derived from host-encoded nitroplast proteins by Coale et al. (2024). MEME-7 and MEME-9 were excluded from every analysis because they overlap the conserved cryptochrome/photolyase functional region and could identify protein-family membership rather than a targeting sequence. The retained motif set comprised MEME-1 through MEME-6, MEME-8, and MEME-10. MAST was rerun against the complete nuclear proteome using ‘-ev 1000000 -remcorr -nohtml -notext’ to retain sequence-level scores for all proteins. A protein was classified as uTP-positive when its MAST sequence E value was ≤10 and at least three distinct retained motifs each had a match with *P* ≤ 1 × 10^-5^ beginning at or beyond 45% of protein length.

N-terminal targeting signals were predicted with TargetP 2.0 using the non-plant model and short-output format (Almagro Armenteros et al., 2019). Proteins assigned a mitochondrial transit peptide (mTP) were classified as mitochondrial-targeting candidates. TargetP signal-peptide (SP) predictions were evaluated with ASAFind 2.0 using the default scoring matrix and thresholds (Gruber et al., 2015, 2025). ASAFind high- and low-confidence predictions were classified as plastid-targeting candidates, whereas the remaining TargetP SP predictions were classified as non-plastid secretory candidates.

Plastid and other secretory classifications were mutually exclusive because ASAFind was applied to TargetP SP sequences; TargetP SP and mTP classes were also mutually exclusive.

### Comparative organelle genome alignment

The *B. bigelowii* FR-21 chloroplast (contig_12873; 107,247 bp) and mitochondrial (ptg001320l; 44,277 bp) sequences described above were compared with haptophyte organelle genomes obtained from GenBank. The main chloroplast comparison comprised *B. bigelowii, Emiliania huxleyi* (NC_007288.1), *Diacronema lutheri* (NC_020371.1), *Chrysochromulina parva* (NC_036937.1), *Tisochrysis lutea* (NC_040291.1), and *Phaeocystis globosa* (NC_021637.1). The expanded chloroplast comparison additionally included *Chrysochromulina sp*. CCMP291 (KJ201907.2), *Isochrysis galbana* (NC_049168.1), and *Phaeocystis antarctica* (PX586075.1). The main mitochondrial comparison comprised *B. bigelowii, C. parva* (NC_036938.1), *Chrysochromulina sp.* CCMP291 (KJ201908.1), *E. huxleyi* (NC_005332.1), *I. galbana* (NC_066487.1), and P. globosa (NC_059693.1). The expanded primary mitochondrial comparison additionally included *P. antarctica* (PX586060.1). Taxa, sequence lengths, and accessions are summarized in Supplementary Dataset S5.

Circular organelle genomes were aligned using progressiveMauve (Darling et al., 2010) with default alignment settings. For visualization, an initial alignment was generated for each organelle dataset and the largest LCB present in every included genome was selected as an anchor. Each sequence was rotated so that the left boundary of its anchor LCB became position 1. Sequences in which the anchor aligned on the reverse strand were reverse-complemented before rotation. progressiveMauve was then rerun on the reoriented sequences, and LCB coordinates, presence, length, order, and orientation were parsed from the resulting XMFA files.

For the simplified main-text panels, separate six-taxon alignments were generated from the rotated sequences, and only LCBs present in all six genomes with a median span of at least 1 kb were displayed (Fig. 3A,B). Supplementary plots retained LCBs with a median span of at least 500 bp that were present in at least four of nine chloroplast genomes or at least two of seven mitochondrial genomes (Supplementary Fig. S4A,B).

Genome bars, LCBs, orientation, and connector ribbons were plotted with a Python script using Matplotlib. Blocks below the genome centerline indicate reverse orientation in the rotated alignment, and ribbons connect homologous blocks between adjacent genomes.

### Functional annotation

Predicted proteins were functionally annotated using complementary orthology- and protein-domain-based approaches. Orthology-based functional annotation was performed using eggNOG-mapper v2.1.13 with the eggNOG v5.0.2 database. Protein sequences were searched against the eggNOG protein database using DIAMOND, with an E-value threshold of 0.001, minimum bit score of 60, minimum sequence identity of 40%, and minimum query and subject coverage of 20%. Taxonomic scope was determined automatically, all ortholog types were considered for annotation transfer, and Gene Ontology annotations were restricted to non-electronic evidence. Orthology assignments were used to obtain functional descriptions, Gene Ontology terms, KEGG Orthology identifiers, KEGG pathways and modules, enzyme commission numbers, and other eggNOG functional annotations.

Protein-domain and family annotations were generated using InterProScan v5.75-106.0 with the default set of available member-database analyses and otherwise default parameters. InterProScan results were exported in tab-delimited format and used to identify conserved protein families, domains, and functional sites.

The Prodigal-predicted nitroplast CDS from contig_13101 were annotated separately using the same eggNOG-mapper workflow. Nitroplast annotations were subsequently incorporated into the compartment-specific annotation framework used for downstream analyses. Nuclear and organellar sequence identifiers retained explicit compartment labels to allow nuclear, nitroplast, chloroplast, and mitochondrial sequences to be distinguished during downstream filtering and analysis. Genome-wide eggNOG annotations for the complete and low-repeat/TE-filtered gene sets, together with organelle identifier crosswalks and sequence-quality flags, are provided in Supplementary Dataset S11.

For functional-overlap comparisons, eggNOG KEGG Orthology identifiers and Pfam terms were deduplicated within each gene set. Figure 2C compares terms in models with at least 50% CDS-repeat overlap with those in models with less than 10% overlap. Figure 2E compares the 166 uTP candidates in the low-repeat set with the 1,685 nitroplast-encoded proteins; Supplementary Fig. S3 additionally includes the comparison using all 604 nuclear uTP candidates. Venn regions count distinct annotation terms rather than proteins.

### Comparative genome and gene-content analysis

Genome size and protein-coding gene number were compared among *B. bigelowii* and publicly available haptophyte genomes, with selected diatom genomes included for broader context. Comparison genome assemblies and annotations were obtained primarily from the DOE Joint Genome Institute (JGI) PhycoCosm database, with corresponding GenBank assembly accessions used where available. Genome sizes were calculated from nuclear genome FASTA sequences where possible, and protein-coding gene counts were obtained from the corresponding genome annotations. Only nuclear genome sequence and nuclear protein-coding genes were included in these comparisons; organelle genomes and organelle genes were excluded. The genome resource, strain, assembly version, source, and accession or PhycoCosm identifier for each taxon are provided in Supplementary Dataset S2.

For comparative phylogenomics, a set of predicted proteomes was assembled for 13 haptophytes: *B. bigelowii, Emiliania huxleyi, Diacronema lutheri, Prymnesium parvum, Phaeocystis cordata, Phaeocystis globosa, Phaeocystis antarctica, Chrysochromulina tobinii, Chrysochromulina parva, Dicrateria rotunda, Isochrysis galbana, Haptolina ericina, and Pavlova sp.* The *P. parvum* UTEX LB 2797 proteome was used as the single representative of *P. parvum* in the species-level OrthoFinder analysis. *Phaeodactylum tricornutum and Thalassiosira pseudonana* were included as diatom outgroups. Full genome and proteome resource information is provided in Supplementary Dataset S2.

Orthogroups and rooted species trees were inferred using OrthoFinder v3.1.3. Sequence similarity searches were performed with DIAMOND (-S diamond), using 24 threads for sequence searches (-t 24) and 24 threads for downstream analysis (-a 24).

Separate analyses were performed using the haptophyte-only dataset and the haptophyte-plus-diatom dataset. The haptophyte-plus-diatom analysis was used to infer the phylogenomic placement of *B. bigelowii* relative to other haptophytes, with *Phaeodactylum tricornutum* and *Thalassiosira pseudonana* included as diatom outgroups (Fig. 1D).

### Comparative analysis of nitrogen metabolism

Nitrogen-metabolism proteins were examined in *B. bigelowii* and the 12 comparison haptophytes. Predicted proteomes from all taxa were functionally annotated using eggNOG-mapper v2.1.13 with the eggNOG v5.0.2 database using the same parameters described above. Candidate homologs were identified from these standardized annotations by searching for a predefined set of nitrogen-metabolism protein families and associated functional annotation terms. Candidate assignments were then manually reviewed to remove unsupported or nonspecific matches and to resolve ambiguous annotations. For *B. bigelowii*, only proteins assigned to the host nuclear genome were considered when evaluating host nitrogen metabolism; nitroplast, chloroplast, mitochondrial, and bacterial proteins were excluded using the compartment-specific sequence identifiers established during genome curation.

The nitrogen gene set represented major pathways of nitrogen acquisition, assimilation, and recycling. Nitrate and nitrite metabolism included nitrate transporters (Nrt), NAR1/FNT-family transporters, nitrate reductase (NR), and nitrite reductase (NirA). Ammonium metabolism included AMT-family ammonium transporters, glutamine synthetase (GS/GlnA), glutamate synthase (GOGAT/GLU), and glutamate dehydrogenase (GDH/GDH2). Urea utilization included DUR3-family urea transporters, other proteins annotated with urea-channel activity where applicable, and urease. Cyanate and nitronate metabolism included cyanate lyase (CynS) and Ncd2-like nitronate monooxygenases.

Arginine metabolism included carbamoyl-phosphate synthase (CPS1), ornithine carbamoyltransferase (OTC), argininosuccinate synthase (ArgG), argininosuccinate lyase (ArgH), and arginase (Arg). Glutaminase and urea amidolyase were evaluated during curation but excluded from the final displayed marker set.

For each taxon and marker family, individual supported protein homologs were counted after manual curation. Curated protein counts were retained in the underlying comparison table, whereas the final nitrogen-metabolism matrix was converted to binary presence or absence for visualization. A marker was scored as present when at least one supported homolog remained after manual review and absent when no supported candidate was retained. The full list of nitrogen metabolism genes is available in Supplementary Dataset S6.

### Flavodoxin candidate identification, uTP-motif screening, and phylogenetic analysis

Flavodoxin homologs were identified in the proteomes of *B. bigelowii* and 12 comparison haptophytes using the eggNOG-mapper annotations described above. Proteins were retained as flavodoxin candidates when they had (i) an eggNOG flavodoxin annotation together with the PFAM Flavodoxin_1 domain, (ii) a K03839 or COG0716 assignment, or (iii) a direct flavodoxin annotation when no PFAM assignment was available. Multi-domain redox proteins and unrelated flavoproteins, including FAD/NAD-binding reductases, sulfite or nitrite reductases, radical-SAM proteins, nitric-oxide synthases, and P450 fusions, were excluded by manual curation. For *B. bigelowii*, candidates were recovered from the complete nuclear eggNOG annotation rather than the conservative low-repeat gene set, preventing exclusion of genes overlapping annotated repeats. This procedure recovered 14 nuclear *B. bigelowii* flavodoxin candidates (10 from the low-repeat set and 4 containing some repeat overlap). Supporting eggNOG annotations and *B. bigelowii* CDS-repeat overlap are provided in Supplementary Dataset S7, full-length protein sequences in Supplementary Dataset S9, and *B. bigelowii* CDS sequences in Supplementary Dataset S10. uTP-associated motif evidence was assigned using the genome-wide MAST procedure described above. The same criterion was applied to *B. bigelowii* and comparison-haptophyte sequences. For *B. bigelowii*, repeat overlap was calculated as the percentage of each coding sequence intersecting the RepeatMasker annotation. Candidate proteins were assigned broad eukaryotic, bacterial, or cyanobacterial categories from their eggNOG orthogroup annotations; these categories summarize sequence similarity and were not treated as direct evidence of evolutionary origin. For phylogenetic analysis, the N-terminal 210 amino acids of each flavodoxin candidate, or the complete sequence when shorter than 210 amino acids, were aligned using MAFFT v7.526 (Katoh C Standley, 2013) with the -auto strategy. A maximum-likelihood tree was inferred using IǪ-TREE v3.1.3 (Wong et al., 2026). ModelFinder selected Ǫ.PFAM+F+G4 as the best-fit amino-acid substitution model according to the Bayesian information criterion (Kalyaanamoorthy et al., 2017; Minh et al., 2021). Branch support was assessed with 1,000 ultrafast bootstrap replicates and 1,000 SH-aLRT replicates; ultrafast bootstrap values ≥70% were displayed. The tree was used to summarize relationships among the candidate flavodoxin sequences. Candidates were classified as eukaryotic or bacterial based on their eggNOG orthogroup assignments.

### Cryptochrome/photolyase candidate identification and phylogenetic analysis

Cryptochrome/photolyase candidates were identified from the *B. bigelowii* nuclear proteome and the same 12 comparison haptophyte proteomes used for the flavodoxin analysis. Proteins were retained when eggNOG assigned COG0415 or KOG0133 together with a ‘DNA_photolyase’ or ‘FAD_binding_7’ domain, or when the direct functional annotation contained photolyase, cryptochrome, or blue-light-photoreceptor terminology. Counts represent predicted protein models. Candidate sequences and annotations were manually reviewed for redundancy and minimum sequence length. uTP status was assigned using the same corrected eight-motif, three-distinct-motif procedure described above. For *B. bigelowii*, CDS-repeat overlap and repeat/TE quality-control classification were appended to each candidate record (Supplementary Dataset S8). Candidate annotations are provided in Supplementary Dataset S8, full-length protein sequences in Supplementary Dataset S9, and available CDS sequences in Supplementary Dataset S10. Terminal motif positions are shown in Supplementary Fig. S7.

Cryptochrome/photolyase candidates at least 150 amino acids long were combined with eight manually reviewed UniProtKB/Swiss-Prot reference proteins representing major cryptochrome and photolyase groups. The resulting 157 haptophyte candidates and eight references were aligned with MAFFT v7.526 (Katoh C Standley, 2013) using the L-INS-i strategy (-localpair -maxiterate 1000). The alignment was restricted to the 489 positions corresponding to residues in the manually reviewed *Synechocystis* sp. PCC 6803 CRY-DASH reference P77967, thereby excluding variable terminal regions including uTP-associated extensions from phylogenetic inference. A maximum-likelihood tree was inferred with IǪ-TREE v3.1.3 under the LG+F+R4 model, with support estimated using 1,000 ultrafast-bootstrap and 1,000 SH-aLRT replicates (Wong et al., 2026). The complete tree was rooted on the class I cyclobutane-pyrimidine-dimer photolyase from *Synechocystis* sp. PCC 6803 (UniProt Ǫ55081). The full 165-tip tree is shown in Supplementary Fig. S8, whereas Fig. 5D displays P77967 and its haptophyte sister lineage. Functional labels indicate phylogenetic similarity to curated references and do not constitute biochemical assignments.

## Supporting information

Supplementary information for: The Braarudosphaera bigelowii genome reveals host remodeling around a nitrogen-fixing organelle

## Notes

### Competing Interest Statement

The authors have declared no competing interest.

### Summary of Updates

THe author list was in incorrect author. T. H. Coale should be the first author, and J. P. Zehr should be the last/senior author.

### Funding

- Simons Foundation

