## Supplementary information for: The Braarudosphaera bigelowii genome reveals host remodeling around a nitrogen-fixing organelle for "The *Braarudosphaera bigelowii* genome reveals cellular remodeling around a nitrogen-fixing organelle"

**Supplementary Figures**

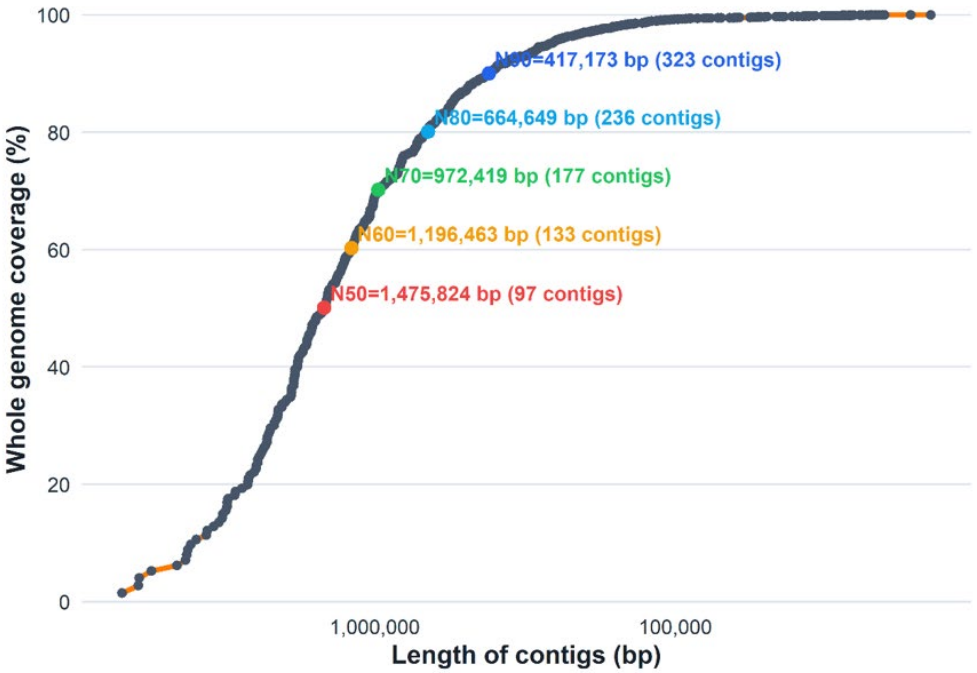


**Figure S1. Contiguity of the B. bigelowii FR21 nuclear assembly.** Contigs are ranked from longest to shortest, and cumulative assembly length is expressed as a percentage of total assembly length. The horizontal axis shows contig length in base pairs on a reversed logarithmic scale. Colored points identify N50, N60, N70, N80 and N90; each label gives the contig length at that threshold and the cumulative number of contigs required to reach it. The N50 is 1,475,824 bp, reached with 97 contigs.

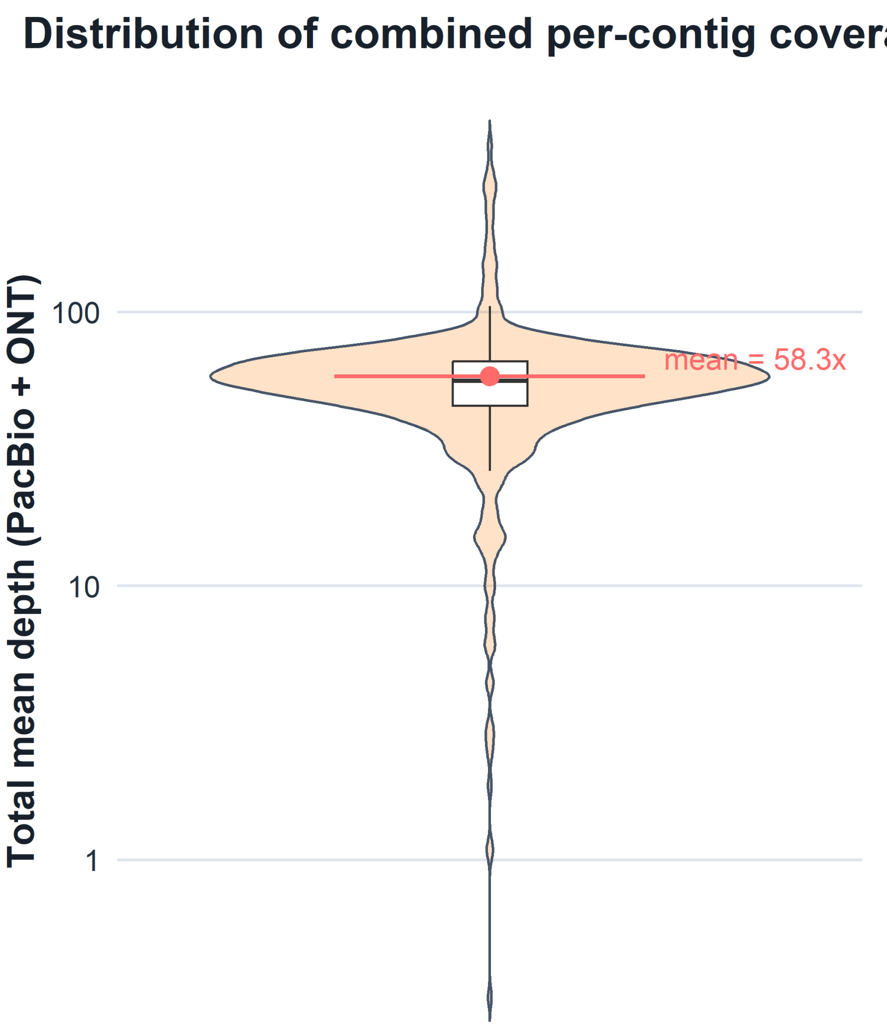


**Figure S2. Combined long-read coverage of the B. bigelowii FR21 nuclear assembly.** PacBio HiFi and Oxford Nanopore reads were mapped independently to the nuclear assembly, and the mean depths for each contig were summed. The violin shows the distribution of combined per-contig mean depth on a logarithmic axis. The inset box shows the median and interquartile range, with whiskers extending to the most extreme values within 1.5 times the interquartile range. The pink point and horizontal line indicate the arithmetic mean across contigs (58.3x), with each contig weighted equally.

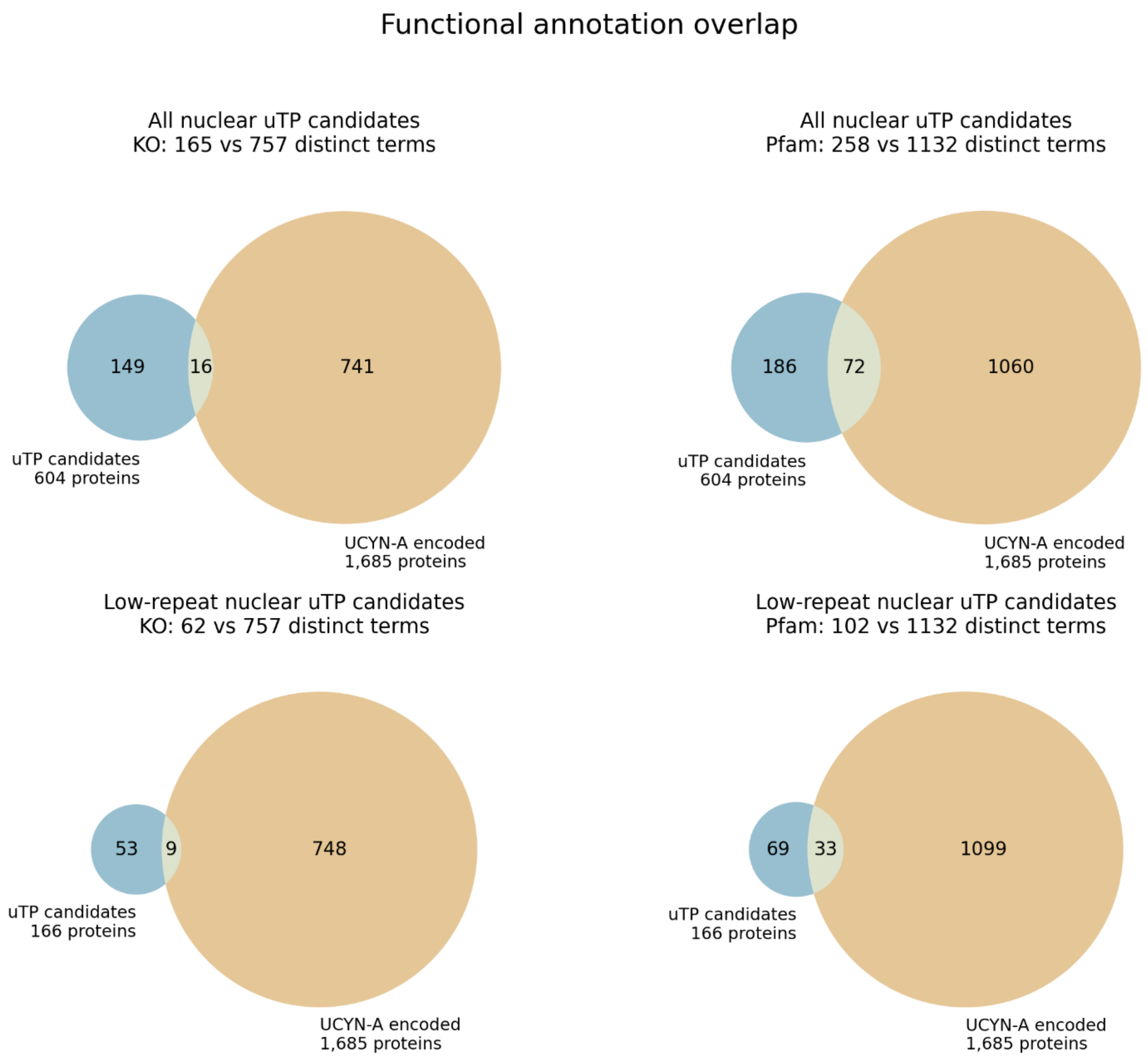


**Figure S3. Expanded functional annotation overlap between nuclear uTP candidates and nitroplast-encoded proteins.** Venn diagrams compare distinct eggNOG KEGG Orthology (KO; left) and Pfam (right) terms in all 604 nuclear uTP candidates (top) and the 166 low-repeat nuclear uTP candidates (bottom) with terms in 1,685 UCYN-A/nitroplast-encoded proteins. Low-repeat is defined as <10% CDS-repeat overlap and no detected TE-domain evidence. Both candidate sets use the three-motif uTP criterion described in Methods. Annotation-overlap counts are provided in Supplementary Dataset S4. Blue denotes nuclear candidates and tan denotes nitroplast-encoded proteins. Numbers within the regions count distinct annotation terms, whereas set labels give protein counts. All candidates share 16 KO and 72 Pfam terms with the nitroplast; curated candidates share 9 KO and 33 Pfam terms. Circle and overlap areas represent term counts within each panel.

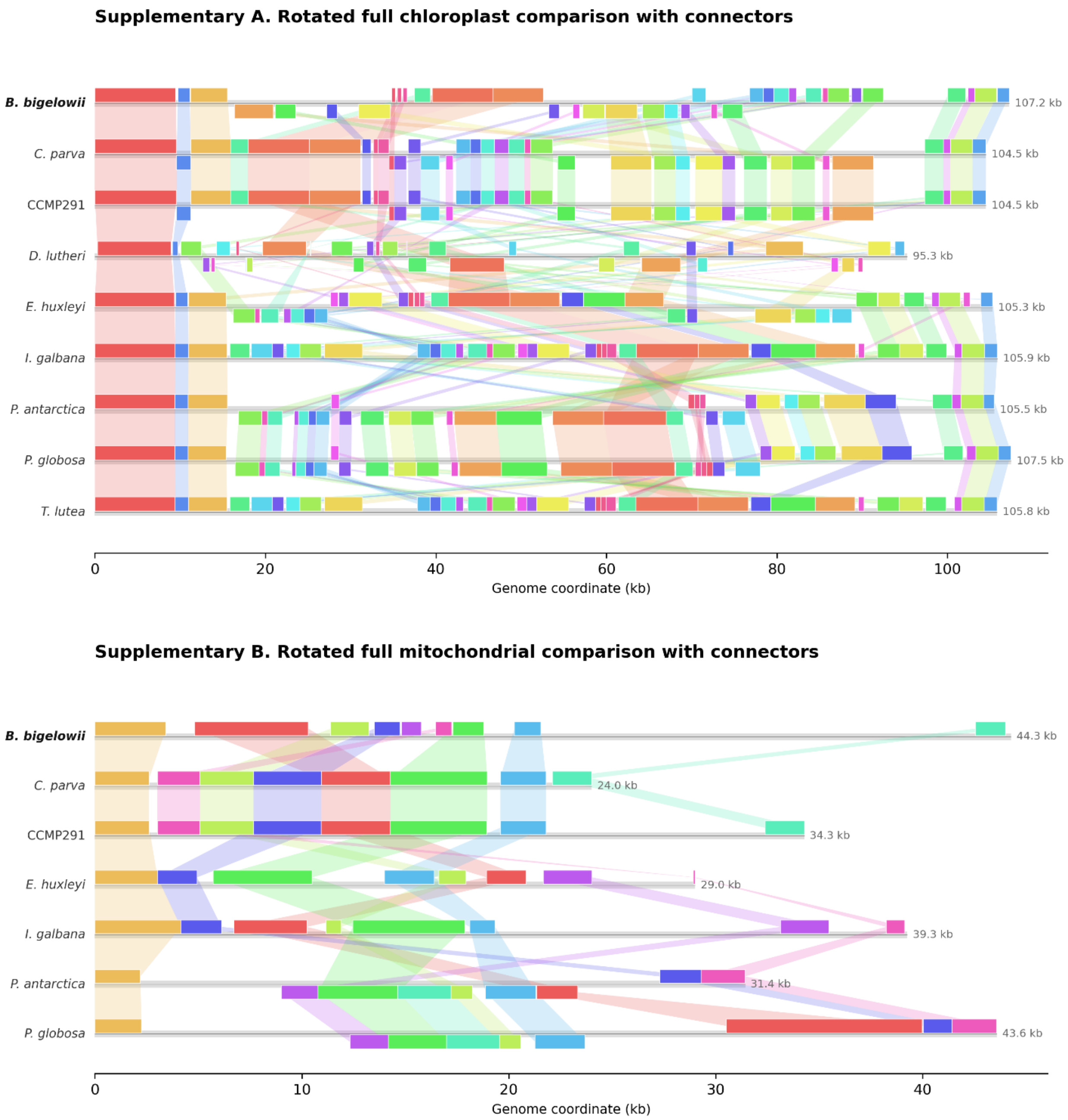


**Figure S4. Expanded comparisons of haptophyte organelle genome organization.** (A) Chloroplast and (B) mitochondrial genomes aligned with progressiveMauve after rotation to a common anchor locally collinear block (LCB). Gray baselines represent complete genome lengths; colored blocks and connecting ribbons identify homologous LCBs within each alignment. Blocks below the baseline are in reverse orientation. Chloroplast blocks have a median span of at least 500 bp and occur in at least four genomes; mitochondrial blocks have a median span of at least 500 bp and occur in at least two genomes. Coordinates are shown in kilobases for the rotated genomes. The corresponding six-taxon comparisons are shown in Fig. 3A,B; genome resources and alignment-summary metadata are provided in Supplementary Dataset S5.

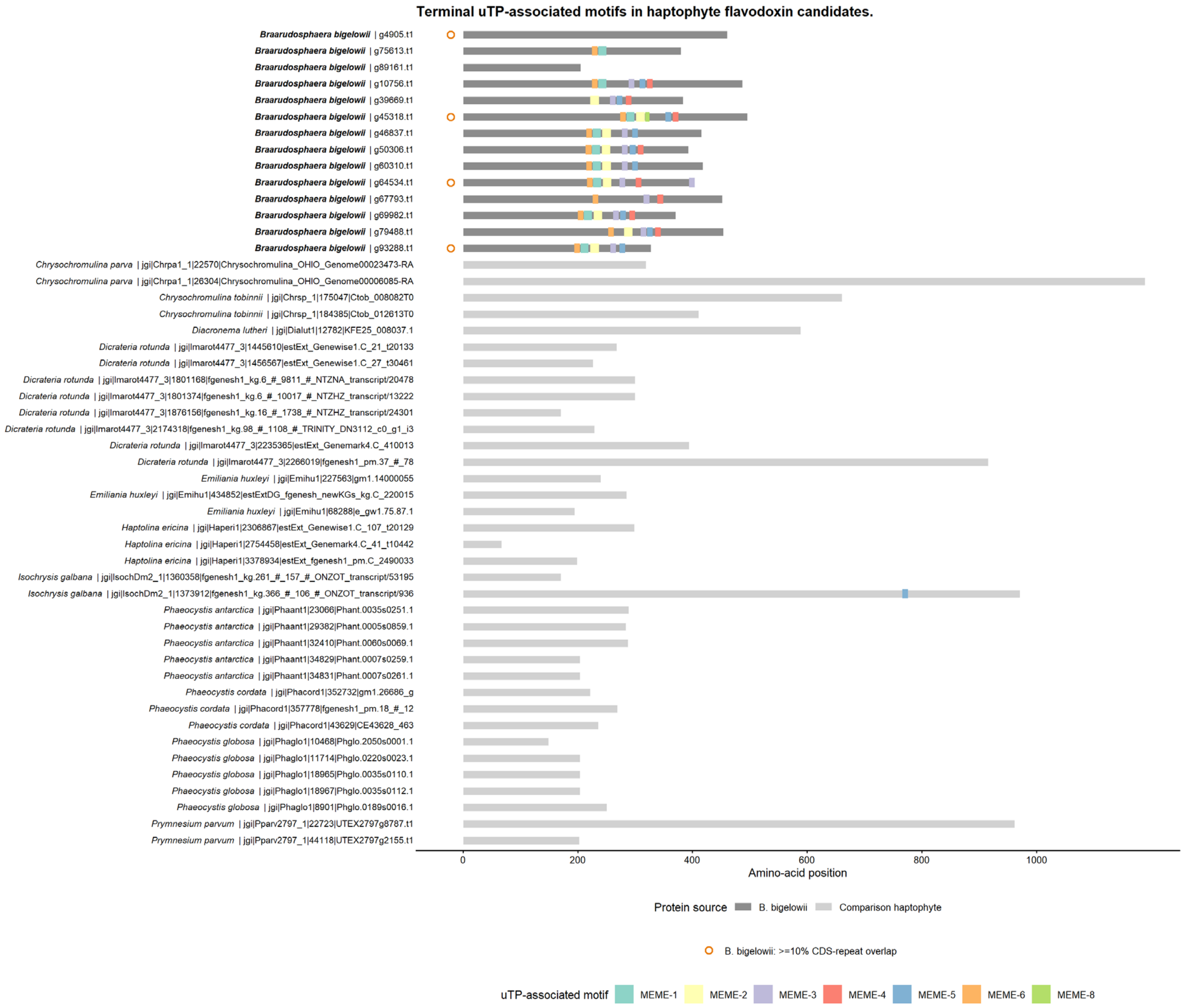


**Figure S5. Terminal uTP-associated motifs in haptophyte flavodoxin candidates.** Horizontal bars represent full-length proteins for the 14 *B. bigelowii* and 36 comparison candidates; bar shading distinguishes the two protein sources. Colored blocks denote individual retained MEME motif matches with MAST P ≤1 × $10^{-5}$ beginning at or beyond 45% of protein length. Positions are in amino acids. Orange circles indicate *B. bigelowii* genes that overlap (≥10% CDS overlap) annotated repetitive regions in the nuclear genome. Full annotations and protein sequences are available in Supplementary Dataset S7 and S9, respectively.

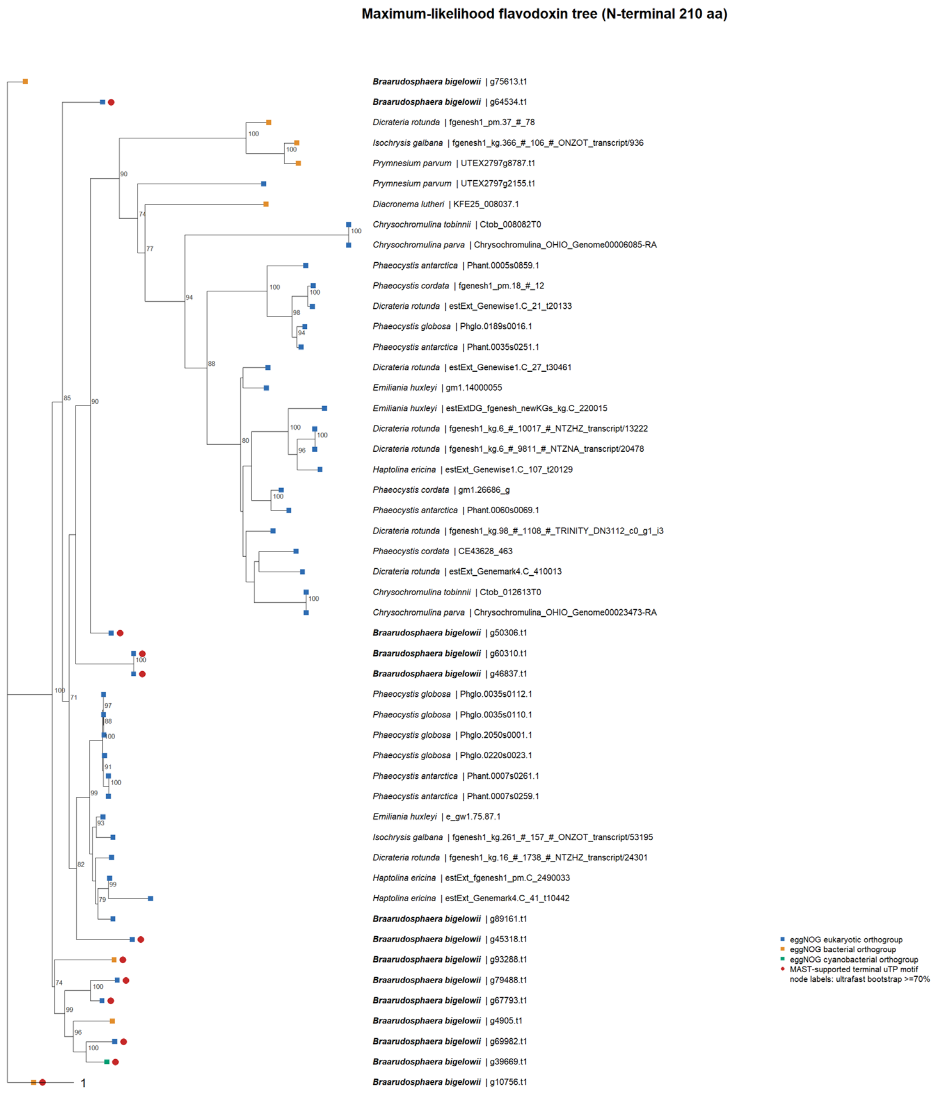


**Figure S6. Expanded maximum-likelihood tree of haptophyte flavodoxin candidates.** The tree includes 14 *B. bigelowii* candidates and 36 candidates from comparison haptophytes. The N-terminal 210 amino acids of each protein, or the complete sequence for shorter proteins, were aligned with MAFFT and analyzed with IQ-TREE under Q.PFAM+F+G4. Node labels show ultrafast bootstrap support ≥70% from 1,000 replicates; the scale bar indicates substitutions per site. Blue, orange and green squares denote eukaryotic, bacterial and cyanobacterial eggNOG orthogroup assignments, respectively. > Red circles identify proteins with a MAST sequence E value ≤10 and at least three distinct retained motifs, each matching at $P \leq 1 \times 10^{-5}$ and beginning at or beyond 45% of protein length. A simplified presentation of the same phylogeny is shown in Fig. 5C; candidate annotations are provided in Supplementary Dataset S7.

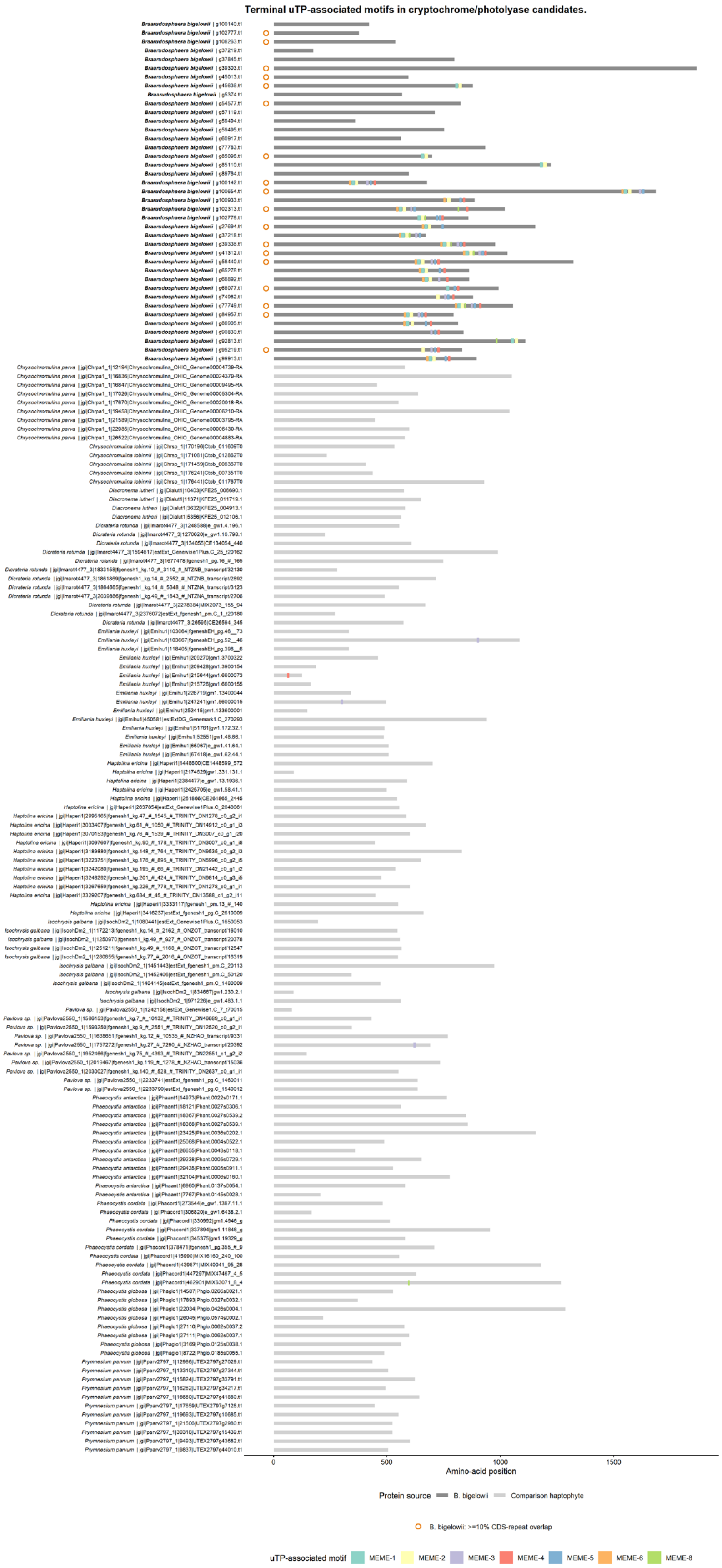
Terminal uTP-associated motifs in cryptochrome/photolyase candidates.
0
500
1000
1500
Amino-acid position
Protein source
B. bigelowii
Comparison haptophyte
B. bigelowii: >=10% CDS-repeat overlap
uTP-associated motif
MEME-1
MEME-2
MEME-3
MEME-4
MEME-5
MEME-6
MEME-8

**Figure S7. Terminal uTP-associated motifs in cryptochrome/photolyase candidates.** Full-length proteins are shown for all 163 candidate models from *B. bigelowii* and 12 comparison haptophytes. Gray bars show protein lengths, and colored blocks show retained MEME motif matches with MAST P $\leq 1 \times 10^{-5}$ beginning at or beyond 45% of protein length. Orange circles indicate *B. bigelowii* genes that overlap (≥10% CDS overlap) annotated repetitive regions in the nuclear genome. Full annotations and protein sequences are available in Supplementary Dataset S8 and S9, respectively.

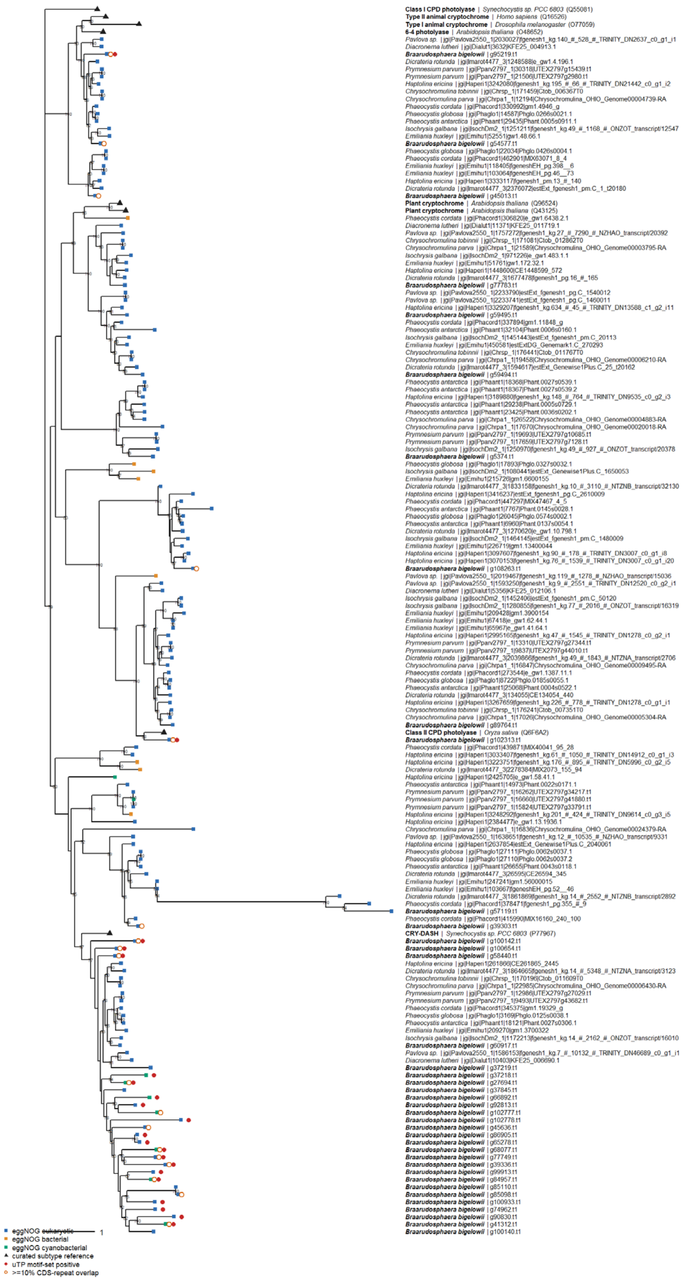

**Figure S8. Expanded cryptochrome/photolyase maximum-likelihood tree.** The tree contains 157 haptophyte candidates ≥150 amino acids long and eight reviewed subtype references. Inference used 489 aligned positions of the photolyase-homology region anchored to *Synechocystis* CRY-DASH P77967, excluding variable terminal regions, including uTP-associated extensions. The tree is rooted on *Synechocystis sp*. PCC 6803 class I CPD photolyase Q55081 for display. Node labels show ultrafast bootstrap support

≥70%; branch lengths are substitutions per site. Blue, orange and green squares denote eukaryotic, bacterial and cyanobacterial eggNOG assignments; black triangles identify cryptochrome/photolyase subtype references. Red circles mark proteins meeting the uTP criterion defined in Methods and the Figure S6 caption; open orange circles mark *B. bigelowii* models with ≥10% CDS overlap with annotated repeats. The CRY-DASH-associated subtree is shown in Fig. 5D; candidate annotations are provided in Supplementary Dataset S8.

**List of Supplementary Datasets:**

Supplementary Dataset S1: Supplementary_Dataset_S1_contig_classification.xlsx

Supplementary Dataset S2: Supplementary_Dataset_S2_comparative_genome_resources.xlsx

Supplementary Dataset S3: Supplementary_Dataset_S3_repeats_and_gene_filtering.xlsx

Supplementary Dataset S4: Supplementary_Dataset_S4_targeting_and_UCYN_A_overlap.xlsx

Supplementary Dataset S5: Supplementary_Dataset_S5_organelle_comparisons.xlsx

Supplementary Dataset S6: Supplementary_Dataset_S6_nitrogen_metabolism.xlsx

Supplementary Dataset S7: Supplementary_Dataset_S7_flavodoxins.xlsx

Supplementary Dataset S8: Supplementary_Dataset_S8_cryptochromes_photolyases.xlsx

Supplementary Dataset S9: Supplementary_Dataset_S9_protein_sequences.fasta

Supplementary Dataset S10: Supplementary_Dataset_S10_CDS_sequences.fasta
